# Spatial distribution of metabolites in primate retina and its relevance to studies of human metabolic retinal disorders

**DOI:** 10.1101/2022.06.13.496007

**Authors:** Roberto Bonelli, Brendan R E Ansell, Sasha M Woods, Sarah Lockwood, Paul N Bishop, Kamron N Khan, Melanie Bahlo, Marcus Fruttiger

**Affiliations:** Population Health & Immunity Division, WEHI, 1G Royal Parade, Parkville, VIC 3052, Australia; Department of Medical Biology, The University of Melbourne, Melbourne, VIC 3010, Australia; UCL Institute of Ophthalmology, University College London, 11-43 Bath St, London EC1V 9EL, UK; UC Davis, CA National Primate Research Centre, Davis, CA 95616, USA; Division of Evolution, Infection and Genomics, School of Biological Sciences, Faculty of Biology, Medicine, and Health, University of Manchester, Manchester, M13 9PT, United Kingdom; Manchester Royal Eye Hospital, Manchester University NHS Foundation Trust, Manchester Academic Health Science Centre, Manchester, M13 9WL, United Kingdom; The Leeds Teaching Hospitals NHS Trust, St. James’s Hospital, Leeds LS9 7TF, UK

## Abstract

The primate retina has evolved regional specialisations for specific visual functions. The macula is specialised towards high acuity vision and is an area that contains an increased density of cone photoreceptors and signal processing neurons. Different regions in the retina display unique susceptibility to pathology, with many retinal diseases primarily affecting the macula. To better understand the properties of different retinal areas we conducted an untargeted metabolomics analysis on full thickness punches from three different regions (macula, temporal peri-macula and periphery) of primate retina. Half of all metabolites identified showed differential abundance in at least one comparison between the three regions. The unique metabolic phenotype of different retinal regions is likely due to the differential distribution of different cell types in these regions reflecting the specific metabolic requirements of each cell type. Furthermore, mapping metabolomics results from macula-specific eye diseases onto the region-specific distributions of healthy primate retina revealed differential abundance defining systemic metabolic dysregulations that were region specific, highlighting how our results may help to better understand the pathobiology of retinal diseases with region specificity.

## Introduction

Systemic metabolic dysregulation can cause pathology in the retina with diabetic retinopathy being a prime example of this. It has been recognised that diabetes long-term metabolic dysregulations can lead to complications in the retina and vision loss with hyperglycaemia believed to be one of the main disease drivers. More recently, several studies have aimed to investigate potential links between other retinal disorders and systemic metabolic changes. For example, metabolomic studies performed on serum from Age-related Macular Degeneration (AMD) have identified associations between dysregulations of lipids as well as amino acids with AMD disease status or severity ^1–4^. Similarly, metabolomic profiling of Macular telangiectasia type 2 (MacTel) patients identified serum levels of serine, and sphingolipids as an important MacTel risk factor ^5–10^. However, it is not clear whether the systemic manifestations (i.e. changes in the serum) are causally related to retinal pathology in AMD or MacTel, or whether they are just indicators of underlying disturbances affecting both retina and peripheral blood.

Complexity is also introduced when retinal diseases do not affect the tissue uniformly due to its spatial structural variation. High acuity vision depends on a region called the macula, which has a peak density of cone photoreceptors as well as retinal ganglion cells and is thicker than the peripheral retina, which is rich in rod photoreceptors. This differential distribution of cells across the retina is reflected by different transcriptional profiles in different retinal regions ^11–17^. However, spatially differential gene expression data presented, so far, limited success explaining why, for example, the macula is particularly affected by diseases such as AMD or MacTel. Furthermore, it is not clear how changes in metabolism, mentioned above, may have a differential impact on retinal diseases. It is also not clear how differential cell distribution impacts metabolite levels in different areas of the retina.

Understanding the spatial distribution of metabolites in the retina, and how that relates to different retinal cell types may therefore be useful to understand how systemic risk factors might affect specific cells in the retina and in particular the macula. This study presents an untargeted metabolomics analysis performed on primates measuring the main metabolic profiles of the different regions of the retina. We identify metabolites and metabolic pathways that differentiate the macula from more peripheral regions and investigate the relationship between these findings and the distribution of different cell types in the retina. Lastly, using our results, we investigate how systemic metabolic risk factors found in both AMD and MacTel relate to the specific metabolic characteristics of the macula.

## Results

### Differential metabolite abundance across the primate retina

To map the distribution of metabolites across the primate retina, samples from three different retinal areas were collected from eleven primate (*Macaca fascicularis)* postmortem eyes and analysed by untargeted metabolomics mass spectrometry analysis (Metabolon Inc.). The raw data (peak areas) underwent log transformation, quality and low abundance filtering, normalisation and missing value imputation. One sample was discarded from further analyses as it was confirmed to be incorrectly collected when checking positive controls (Materials and Methods). The final dataset consisted of 32 retinal samples from six primates. Three females and three males were included in this study with an average age of 2.1 years. For each sample, 371 metabolite abundances were retained. Liner modelling correcting for several covariates was performed as described in Materials and Methods to detect metabolites with differential abundance between retinal areas. We found a total of 197 (53%) metabolites whose abundance was significantly different in at least one statistical contrast (**Table S1**). The number and intersection of differentially abundant metabolites between retinal areas are presented in **Figure S1**. The largest number of differences was found between the macula and periphery (190), the two regions with the largest spatial distance in the retina. We did however find more differentially abundant metabolites between the macula and temporal areas (97), than between the temporal and peripheral areas (53).

Comparing the abundance of individual metabolites in the macula versus periphery (**Figure S2, Table S1**), the most significantly enriched metabolites were carotene diol 1 and 3, corresponding to the macular pigments lutein and zeaxanthin. These two diet-derived xenobiotics are known to be enriched in the macula of primates and were used in this study to validate the correct dissection of the macular samples (Materials and Methods). Of note, ergothioneine and beta-guanidinopropanoate, also diet-derived, were the third and fourth most significantly enriched metabolites, as well as xenobiotics, in the macula. In contrast, the positive control euthanasia drug, pentobartbital, was equally distributed across the retina (**Table S1**). The most enriched endogenous metabolites in the macula were N-acetyl-3-methylhistidine, N-acetyl-aspartyl-glutamate (NAAG) and 2-hydroxyglutarate. The NAA precursors N-acetylaspartate (NAA) and aspartate were also significantly enriched metabolites (rank 15 and 25) (**Table S1**). The most depleted non-lipid metabolites in the macula were putrescine and taurine (rank 2 and 4, **Table S1**).

To better understand which aspects of the metabolism differ between the retinal periphery and the macula, metabolites were grouped according to metabolic pathways and biochemical families and tested for directional enrichment (**Figure 1, Table S1)**. The pathway with the most significant enrichment in the macula was the alanine/aspartate pathway, followed by the tricarboxylic acid (TCA) cycle. Conversely, the macula was relatively depleted of phosphatidylethanolamine, ceramide, sphingomyelin, diacylglycerol, polyamine and benzoate (in order of statistical significance). Overall, there was a similar number of metabolites enriched (99) versus metabolites depleted in the macula (98), but these differences were contributed to by metabolites from different classes. Overall, lipids were depleted in the macula and non-lipid metabolites were enriched, as displayed in **Figure 1, S2)**. Furthermore, four of the six significantly depleted metabolic groups were lipids (note, the bottom four rows in **Figure 1** are primarily depleted).

**Figure 1:**
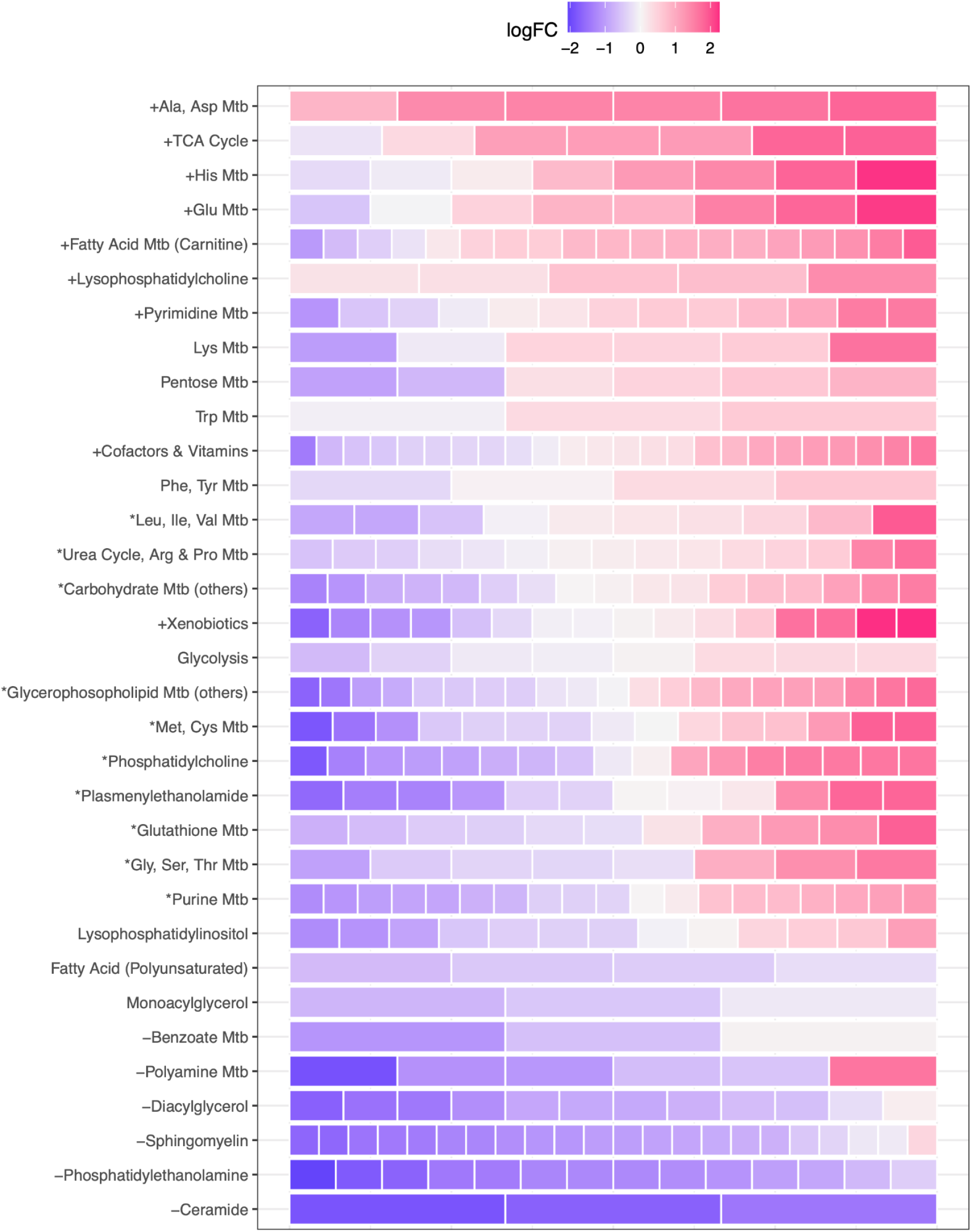
Differential abundance of metabolites across retinal space. Relative abundance of N metabolites (filled rectangles, or ‘bricks’) grouped by biochemical family/pathway (y axis) and ordered by greater family-wise abundance in the macula (top) to the periphery (bottom). The number of bricks indicates the number of metabolites in the pathway available for this study. The colour of each brick represents the log-fold change of that metabolite between macula and periphery. Positive log-fold change (reds) indicates that the metabolite is more abundant in the macula while negative values (blues) indicate that the metabolite is more abundant in the periphery. + denotes family is significantly enriched in the macula (mainly red); - denotes family is significantly depleted in the macula (mainly blue); * denotes family is more differentially abundant in both directions than would be expected by chance (‘mixed directionality’/signal mixture effect).

Plotting the distribution of metabolic pathways and biochemical families in the three sampled regions (macula, temporal and peripheral) further illustrates the relative lack of ceramide, phosphatidylethanolamine, sphingomyelin and diacylglycerol in the macula compared to the periphery (**Figure 2)**. Of note, phosphatidylcholine and plasmenylethanolamine were much more evenly distributed, and carnitine-related fatty acid metabolism and lysophosphatidylcholine were the only lipid groups enriched in the macula.

**Figure 2:**
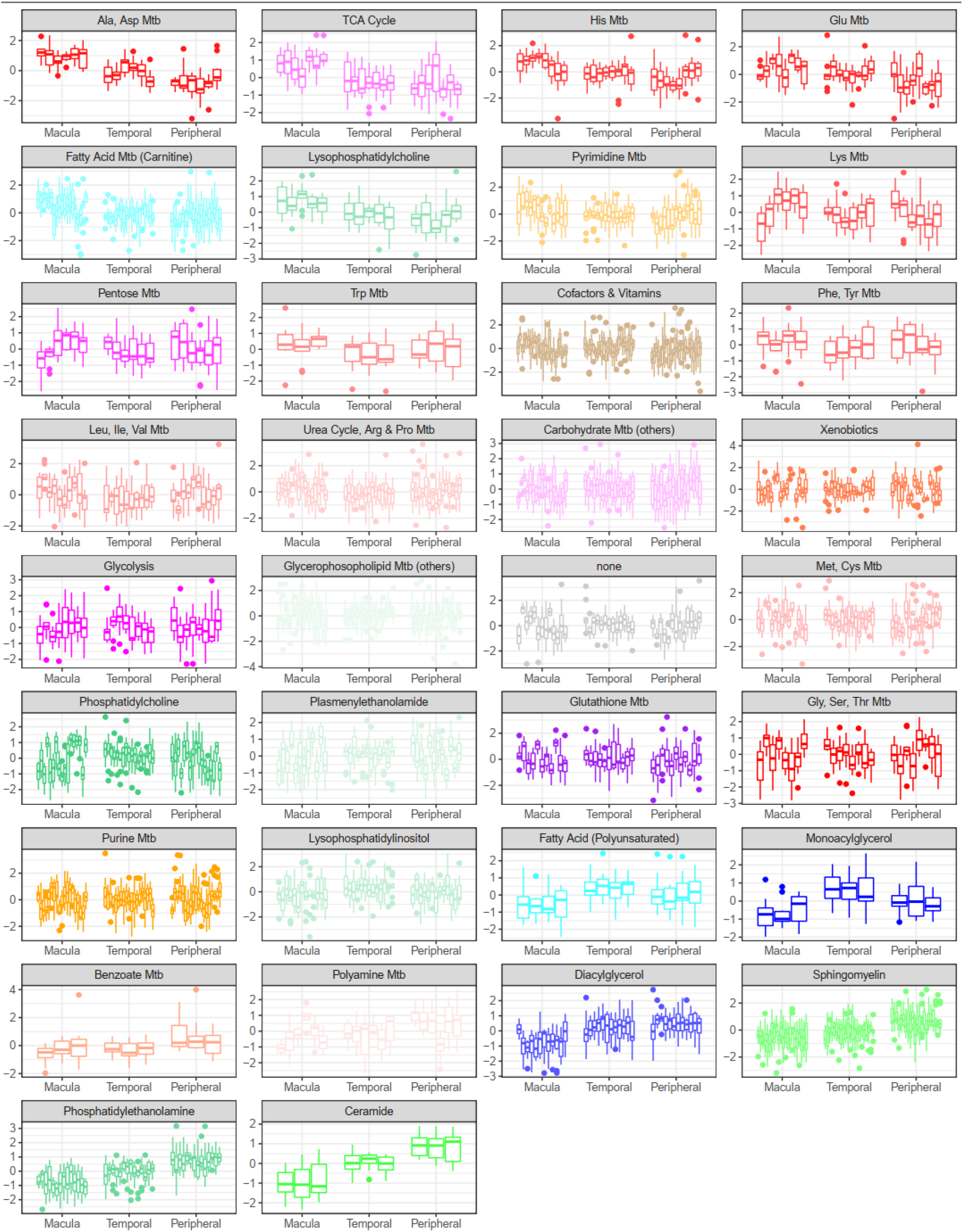
Metabolic distribution of biochemical families between retinal areas. In this image, every boxplot represents the covariate-corrected distribution of one metabolite whose abundance was significantly different for any of the three contrasts. Metabolites are divided into biochemical families which are sorted according to greater family-wise abundance in the macula (top) or in the periphery (bottom).

In addition, we tested for metabolic pathways enriched with differentially abundant metabolites represented in both directions (‘mixed directionality enrichment’). Pathways with significant enrichment/depletion in the macula were the leucine, methionine, glycine/serine, urea cycle, glutathione, purine, phosphatidylcholine and the plasmenylethanolamine metabolite classes (**Figure 1, Table S1**). The individual differentially-abundant metabolites that constitute these mixed directionality results (pathways labelled with a star in **Figure 1**) are displayed in **Figure 3**. In contrast to the other lipid groups, the number of different phosphatidylcholines (dark green in **Figure 3**) was similar in the enriched and depleted categories. However, the unsaturated forms were typically enriched, and the saturated ones tended to be depleted in the macula. Also of note were the purine family metabolites, wherein phosphorylated purines (ADP, AMP, GDP and IMP) were enriched in the macula, and their non-phosphorylated counterparts (adenosine, guanosine and inosine) were depleted.

**Figure 3:**
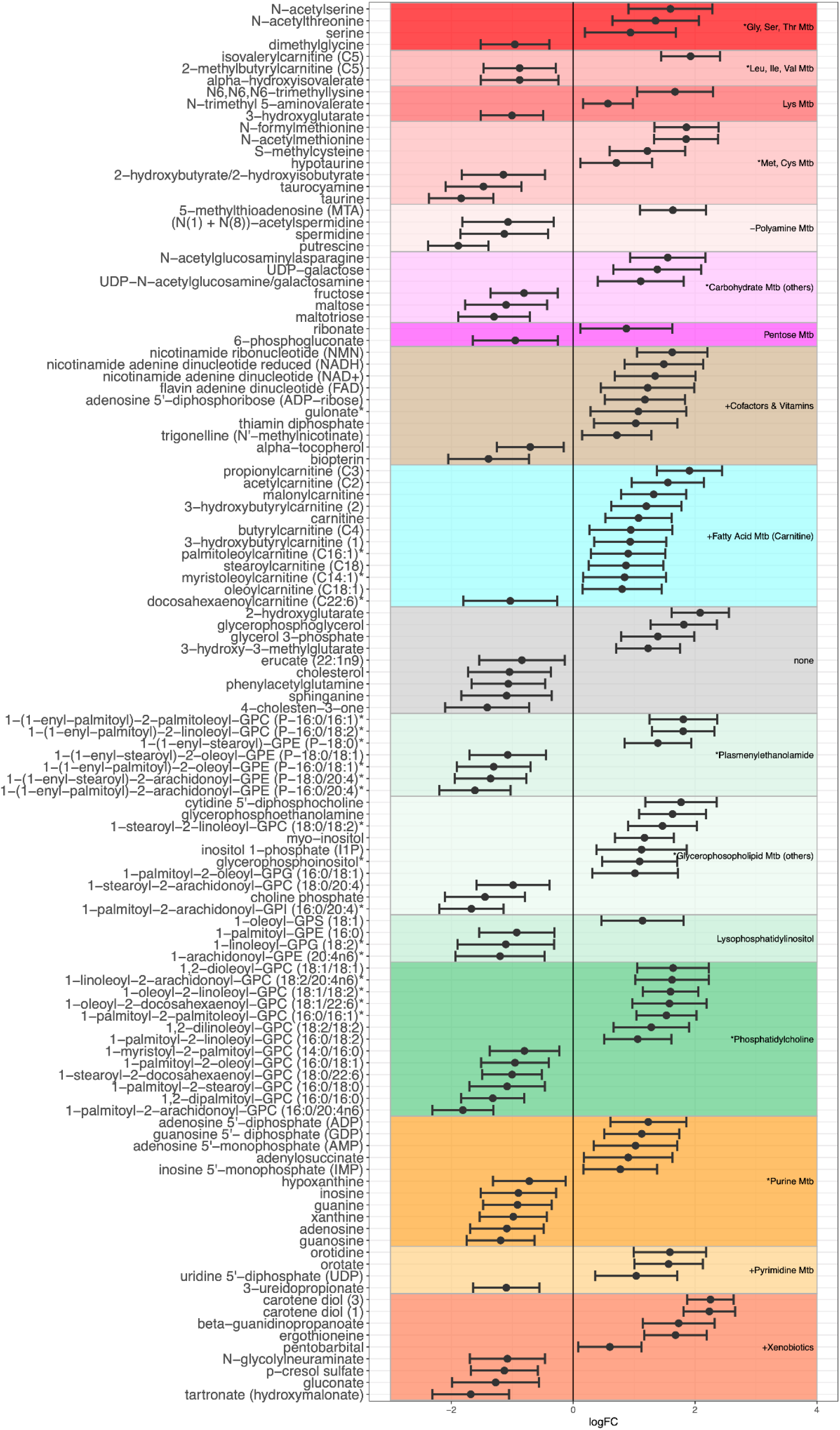
Log-fold changes and 95% confidence interval of metabolite families/pathways with mixed abundances between the macula and periphery. Only metabolites which are individually significantly differentially abundant are displayed. Positive log-fold changes values in this figure indicate that the metabolite abundance was higher in the macula compared to the periphery and *vice versa*. Metabolites are grouped and coloured by their respective biochemical families/biological pathways. + denotes family is significantly enriched in the macula (mainly red); - denotes family is significantly depleted in the macula (mainly blue); * denotes family is more differentially abundant in both directions than would be expected by chance (signal mixture effect). Families with significant abundance entirely in the macula or periphery are omitted for clarity (provided in **Figures S4-6**).

Abundance differences of each metabolite in the macula and peripheral regions are displayed in **Figures S2, S3, and S4**. To explore the distribution characteristics of metabolites across all three sampled retinal areas, we grouped metabolites according to differential abundance patterns that progressed laterally from the macula to the temporal region to the periphery. We defined ten major pattern clusters (**Table 1, Figure S5, Figure S6**), the largest of which contained metabolites that progressively increased or decreased according to proximity to the retinal centre. A total of 173 metabolites showed the greatest abundance differences at the extremes of the macula>temporal>peripheral axis, with only seven showing significant differences in the temporal retina relative to the other regions.

**Table 1.** Abundance pattern cluster descriptions.

| Cluster Name | Description | Significant Pathways |
| --- | --- | --- |
| Steep Enrichment | Metabolites whose abundance very quickly increases when moving from the periphery to the macula of the retina. Differences are consistent in direction and significant between all three areas. | Alanine and Aspartate Metabolism; Histidine Metabolism; Methionine, Cysteine, SAM and Taurine Metabolism |
| Shallow Enrichment | Metabolites whose abundance slowly increases when moving from the periphery to the macula of the retina. A significant difference is only perceived when comparing macula to periphery. | Glutamate Metabolism; Xenobiotics; Carbohydrate Metabolism (other); Glycine, Serine and Threonine Metabolism |
| Macula Enriched | Metabolites whose abundance is particularly elevated only in the macula. | TCA Cycle; Fatty Acid Mtb (Carnitine); Lysophospholipid; Lysine Metabolism |
| Periphery Depleted | Metabolites whose abundance is particularly depleted only in the periphery. | none |
| Steep Depletion | Metabolites whose abundance very quickly decreases when moving from the periphery to the macula of the retina. | Phosphatidylethanolamine (PE); Ceramides; Polyamine Metabolism |
| Shallow Depletion | Metabolites whose abundance slowly decreases when moving from the periphery to the macula of the retina. A significant difference is only perceived when comparing macula to periphery. | none |
| Periphery Enriched | Metabolites whose abundance is particularly elevated only in the periphery. | Sphingomyelin; Benzoate Metabolism |
| Macula Depleted | Metabolites whose abundance is particularly depleted only in the macula. | Diacylglycerol; Phospholipid Metabolism |
| Temporal Enriched | Metabolites whose abundance is particularly elevated only in the temporal region. | Cofactors & Vitamins; Monoacylglycerol; Lysophospholipid |
| Temporal Depleted | Metabolites whose abundance is particularly depleted only in the temporal region. | none |

Similarly, we clustered metabolic families by first deriving principal components (PCs) for each family and then testing for differences in their magnitude between retinal regions. As expected, when these differences in PC abundance were grouped, most pathways/families recapitulated spatial distribution patterns observed using their constituent metabolite abundances (**Table 1, Table S1**).

### The regional retinal context of disease-related metabolites

To place results from our analysis in healthy primate retina into the context of pathology, we mapped our data against serum-based investigations of two retinal disorders that exclusively affect the macula and have known associations with systemic metabolic changes, Macular Telangiectasia type 2 (MacTel) ^5–10^ and Age-related Macular Degeneration (AMD)^1–4^. To this end, we leveraged differential serum metabolite abundance results from our previously published MacTel study ^5^, to compare serum metabolites in patients with MacTel compared to controls. In addition to MacTel, we also recruited a cohort of 205 individuals with AMD and 146 healthy controls. Patients were divided into sub-phenotypes of choroidal neovascularization (CNV), geographic atrophy (GA) and ‘mixed’. Abundances of 763 serum metabolites were compared between each AMD patient subgroup and the controls. Although similar size studies of AMD have already been published ^1–4^, our data and its analysis ensured that the data generation, pre-processing, cleaning and statistical analysis were consistent across datasets. In all statistical contrasts, only four metabolites were identified as being significantly differentially abundant after correcting for multiple testing: tryptophan betaine, heptanoate, 1-linoleolglycerol (18:2) and 1-pentadecanoylglycerol (15:0). All four were depleted in the serum of AMD-CNV patients compared to controls. A further 73 metabolites were differentially abundant in this patient subgroup at the nominal threshold of p < 0.05 (**Supplementary Results** and **Table S2**). We found 36 and 37 nominally significant metabolites when comparing AMD-GA and AMD-Mixed to controls. None of these remained significant after accounting for multiple testing.

To investigate the spatial dimension of patient serum results in the retina, we separated the log fold-change results from the human serum-based studies into three groups according to their specific abundance in the primate macula or periphery, or annotated them as having a non-significant difference (**Figure 4 A-B**). Several of the metabolites that were changed in patient serum were found to be differentially distributed in healthy primates’ retinas. For instance, metabolites of the glycine/serine/threonine metabolism as well as alanine/asparagine metabolism were systemically depleted in the serum of MacTel subjects and were defined by our analyses to be more abundant in the macula compared with periphery or not differentially abundant (**Figure 4A**). The opposite trend was evident for phosphatidylethanolamines, which exhibited lower abundance in the macula than periphery but were enriched in the serum of MacTel patients. Sphingomyelins were significantly more abundant in the macula and depleted in MacTel patient serum. This analysis revealed however some interesting diversification patterns across the methionine and cysteine metabolism whose metabolites depleted in MacTel were mostly macula enriched rather than the non-differentiated ones. In AMD, purine and ceramide metabolism was reduced in patient serum and healthy primate retina (**Figure 4B**).

**Figure 4:**
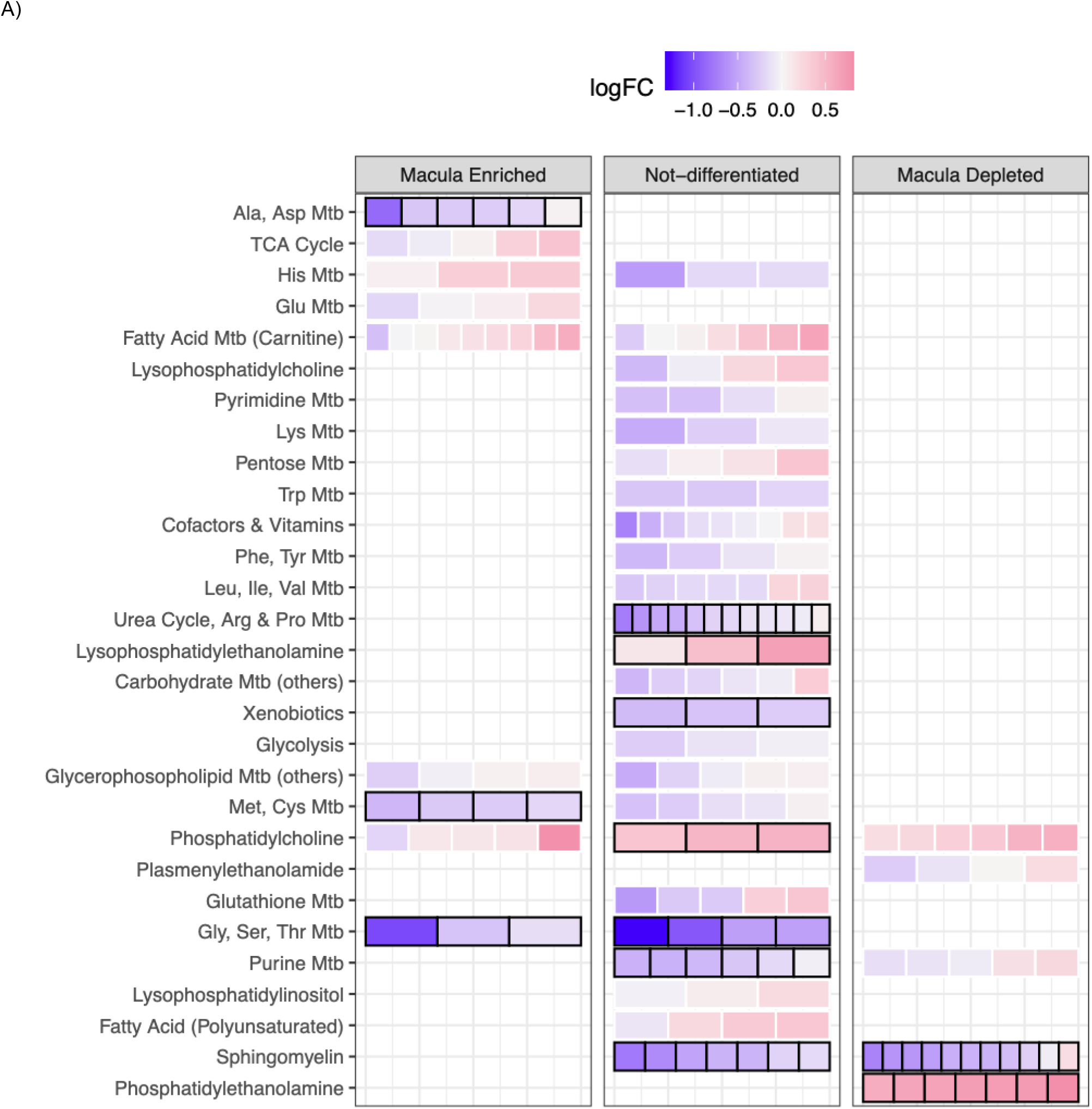
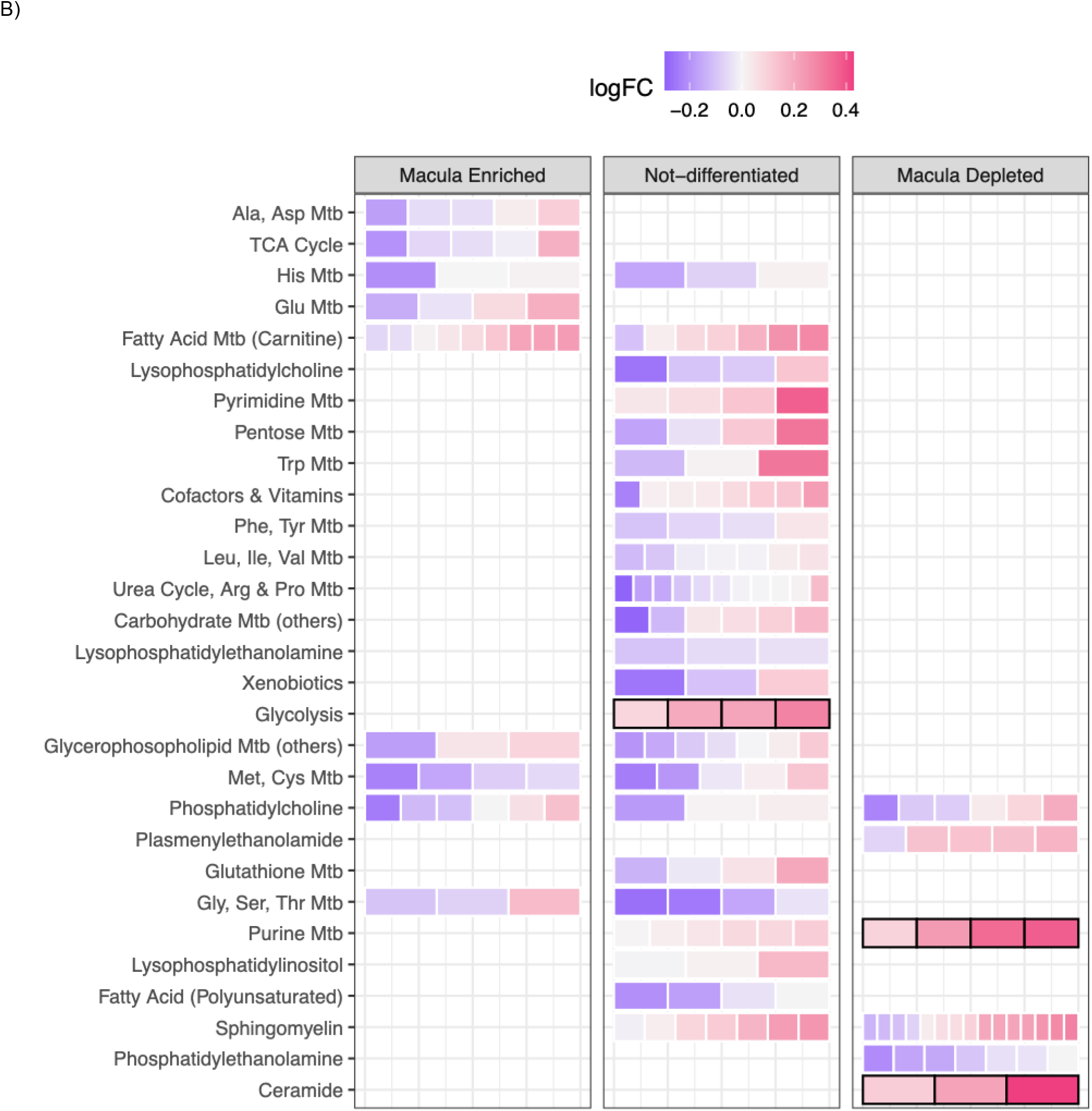
Area Region specific metabolic pathways enrichment results for two retinal diseases: (A) MacTel and (B) AMD. Each row represents a biological pathway that contains metabolites either enriched, not differentiated or depleted in the macula of the primates compared to the periphery. Metabolites are presented as “bricks” in each row. The colour of each metabolite represents the log-fold changes of that metabolite between MacTel cases and controls. High log-fold change indicates that the metabolite is more abundant in the cases compared to controls while negative values indicate a depletion of such metabolite. Sub-pathways significantly enriched (FDR<0.05 for MacTel and nominal p<0.05 for AMD) present a thick black stroke around them. There were no metabolites in the AMD study that achieved FDR<0.05.

## Discussion

In this study, we measured metabolic abundances in three different regions of the primate retina: the central macular, temporal, and peripheral regions. Using a large and untargeted panel of 371 metabolites for 32 retinal samples, our study is the first of its kind elucidating the metabolic profile of retinal regional areas identifying multiple differences. We found a total of almost 200 metabolites whose abundance differed between the three areas, highlighting a clear metabolic separation between the macular region of the retina compared to the outer, temporal or peripheral, areas.

The most enriched metabolites in the macula were identified in our study as carotene diol 1 and 3. These are carotenoids containing an oxygenated carotene backbone and are known as xanthophylls. It is well established that lutein and zeaxanthin are two diet-derived xanthophylls highly enriched in the macula and responsible for the characteristic primate yellow pigment spot ^18^. Since zeaxanthin is relatively more concentrated in the macula compared to lutein ^19^, the distribution pattern observed in our study (**Fig. S7**) indicates that carotene 3 corresponds to zeaxanthin and carotene diol 1 to lutein. Furthermore, the finding also validates that the tissue samples were dissected from the correct retinal region.

The function of these yellow macular pigments has not been firmly established yet, although their natural anti-oxidative properties are proposed to protect the retina from light-induced oxidative damage. In this context, it is interesting that we also identified ergothioneine, a diet-derived, colourless xenobiotic with antioxidative properties ^20^, as the third most enriched xenobiotic in the primate macula – lending further credence to the hypothesis of macula-specific anti-oxidative mechanisms. This concept is further supported by the macular enrichment of glutathione (**Table S1**), which is an endogenously generated antioxidative compound.

The fourth most enriched xenobiotic in the macula, beta-guanidinopropanoate, is known to decrease intracellular creatine and phosphocreatine levels and, in skeletal muscle, to increase fatigue tolerance ^21^. Considering the important role of phosphocreatine in photoreceptor energy metabolism ^22,23^, it is plausible that the specific accumulation of beta-guanidinopropanoate contributes to the fine-tuning of energy usage in the macula.

One of the most striking observations in our study was the reduced abundance of many lipid groups in the macula compared to the periphery. Since many of the lipids detected in our study are membrane constituents, it is likely that this differential distribution is linked to the anatomy of the retina. More specifically, photoreceptors contain more lipids than other cells in the retina because their outer segments consist of densely stacked disc-shaped membranes. The thickness of the outer segment layer is more or less uniform across the entire retina, whilst the inner retina (composed of four major neuronal subtypes) approximately doubles in thickness in the macula compared to the periphery. As all samples in this study (full-thickness retinal punches) were normalised, this roughly halves the relative contribution of outer segments (from rods and cones) in the macular sample. Furthermore, the molar ratio of phosphatidylethanolamine to phosphatidylcholine in disk membranes is much higher (1:1) than in normal plasma membranes (1:6) ^24^, which explains the relatively low abundance of phosphatidylethanolamine in the macula, compared to phosphatidylcholine. By extension, it is plausible that the pronounced relative reduction of sphingomyelin and ceramide abundance in the macula is also based on enrichment of these lipids in photoreceptor outer segments, although this has yet to be validated.

The second-most depleted non-lipid compound in the macula was taurine, which is known to be highly enriched in the retina, more so than any other bodily tissue ^25^. Photoreceptors accumulate taurine and critically depend on it for long term survival ^25–27^. Thus, as in the case of phosphatidylethanolamine described above, the relative depletion of taurine in the macula can be attributed to its known enrichment in photoreceptors. Consequently, we may speculate that putrescine (the most depleted macula non-lipid metabolite) and the polyamine family metabolites (the most depleted macula non-lipid metabolic group) are overrepresented in photoreceptors. However, supporting evidence for this ^28^ is scarce. The spatial distribution of cones and rods might also contribute to the differential abundance of the phosphorylated versus non-phosphorylated purines such as GDP/GTP and guanosine as these metabolites are important components of the visual signalling cascade and might reflect activity differences between cones versus rods in the light-adapted retina.

The most enriched metabolite in the macula was NAAG, which is the third-most-prevalent neurotransmitter in the mammalian nervous system ^29^ after glutamate and taurine. Although the physiological function of NAAG in the retina is not well understood, its synthesis in and release from retinal ganglion cells in mammals and birds is well established by immunohistochemistry ^30,31^ and by the enrichment of the NAAG producing enzymes (N-acetylaspartate synthetase and N-acetylaspartylglutamate synthase) visible in published single-cell transcription profiles ^32^. This suggests that the increased abundance of NAAG is due to the much higher numbers of retinal ganglion cells in the macula versus the periphery. These results show that future studies that allow single cell metabolomics are needed. These will be able to identify cell type specific metabolite repertoires and will alleviate the problem of unknown cell type proportion contributions to these results ^33^.

Considering the important roles specific metabolites play in cytoarchitecture and cellular function it is arguably not surprising that we found in this study over half of all identified metabolites were differentially distributed in at least one comparison. This appears to apply in particular to metabolites associated with retinal ganglion cells, rods and cones, which display the most extreme spatial distribution gradients across the retina. In contrast, we would expect metabolites that are linked to cell types with a more even spread, such as Müller cells or bipolar cells, to show less pronounced abundance differences in our three sampling locations. An illustrative example of this may be gamma-aminobutyrate (GABA) and glutamine which are both evenly distributed (**Table S1**) and which are part of dominant metabolic pathways in Müller cells ^34^.

To explore the relevance of regional metabolic phenotypes in the retina in the context of disease, we explored the potential overlap between our primate retina dataset and serum metabolomics data from patients with retinal eye diseases. We chose to focus on MacTel and AMD because both of these diseases are associated with systemic metabolic changes and both manifest themselves region-specifically in the macula. The retina and the serum studies were all based on the same metabolomics platform (Metabolon) and the bioinformatic pre-processing of all datasets was uniform, enabling meaningful comparisons whereas other studies often have this as a confounder. This approach revealed that the metabolic pathways that were changed in the serum of MacTel patients and that also displayed a region-specific distribution in the healthy retina were in the alanine/asparagine metabolism, parts of glycine/serine metabolism, the phosphatidylethanolamine metabolism and the sphingomyelin metabolism. In AMD, we found that the purine and ceramide metabolic pathways showed retinal region-specificity (in the healthy retina) as well as changes in patient serum.

Although the overlaps between healthy primate retina and human patient serum do not establish causality or imply a functional role in pathobiological mechanisms, they are likely to aid better understanding on why diseases such as MacTel and AMD primarily affect the macula.

## Data Availability

Primate retinal metabolomic data and human plasma metabolomic data are available on request.

## Acknowledgements

We thank Prof Tony Moore and Prof John Yates for their help with sample and data collection from AMD patients. We thank Dr Valentina Cipriani for assisting with matching the serum donor demographics. Finally, we thank patients and their families for their involvement in this study.

## Author Contributions

Conceptualization: MF

Methodology: SMW RB MF

Software: RB

Formal analysis: RB BREA MF

Investigation: RB BREA MB MF

Data collection: SMW PNB KNK

Resources: RB SMW SL MF

Data Curation: SMW SL

Writing - Original Draft: RB BREA MF

Writing - Review & Editing: RB BREA MB MF

Visualization: RB

Supervision: MB MF

Project administration: BREA MF

Funding acquisition: MB MF

## Materials and Methods

### Sample collection and processing

#### Macaque retina collection and processing

Metabolic abundances were measured in the retinas of six healthy adult primates (*Macaca fascicularis*). For all animals but one, both eyes were included in the study for a total of 11 retinas. Fresh neural retinal tissue in the light-adapted state was dissected 10-20 minutes after euthanasia from three different eccentricities from the fovea. The yellow macular pigment spot (macula lutea) was clearly visible in the dissected retinas and used to locate the fovea, from where a piece of retinal tissue was cut out (centred on the fovea). The diameter of the sample was equivalent to the distance between the fovea and the temporal edge of the optic disc. A second, adjacent sample of the same size was taken temporally to the foveal sample. A third sample (same size) was taken further temporally in the peripheral retina, resulting in a total of 33 samples. The primates presented an average age of 2.1 years (sd=0.94) and were divided into 3 females and 3 males. Samples were collected and immediately flash frozen on 3 separate dates and then submitted to Metabolon Inc. (Durham, USA) for mass spectrometry analysis. Briefly, this involved analysis of five fractions per sample: two for analysis by two separate reverse phase (RP)/UPLC-MS/MS methods with positive ion mode electrospray ionization (ESI); one for analysis by RP/UPLC-MS/MS with negative ion mode ESI; and one for analysis by HILIC/UPLC-MS/MS with negative ion mode ESI. Raw data was extracted, peak-identified and QC processed using Metabolon’s hardware and software. Compounds were identified by comparison to library entries of purified standards and peaks were quantified using the area-under-the-curve technique, providing relative abundances of 435 metabolites.

#### Human blood collection and processing

To assess the utility of our results when exploring systemic metabolic profiles of retinal disorders, we collected metabolic abundances from plasma of 351 participants. Of these 205 were AMD patients, while the rest were age-matched controls without AMD. AMD patients were divided into three sub-disease categories according to their retinal diagnosis, patients with choroidal neovascularization (CNV), geographic atrophy (GA) and patients with CNV and GA (mixed). This data consisted of 127 CNV, 45 GA and 33 Mixed AMD patients. The metabolic measurements were processed in 12 batches. The participants presented an average age of 78.2 years (sd=7.31) with 54% females and 46% males. For each sample, we received abundances for 1,403 metabolites, 431 of which were discarded *a priori* given that they were not defined as specific metabolites by Metabolon Inc.. Metabolic missingness rate was 14% (sd = 27%) among all samples with a similar missingness rate between healthy individuals (13.8%) and all the AMD cases (CNV 13.7%, GA 13.5%, Mixed 14.4%). Missingness varied across metabolites (**Table S2**).

### Data pre-processing and imputation

Given their strongly skewed distribution, metabolic abundances were log-transformed to achieve distribution symmetry. Metabolic abundances were normalised by dividing the global median log-abundance of area/batch combination. To validate that samples were dissected from the correct locations in the retina we monitored the abundance of the metabolites carotene diol 1 and 3. Although Metabolon has not yet directly assigned the two carotenoids to specific structures, carotene diol 1 and 3 almost certainly represent the macular pigments lutein and zeaxanthin, respectively. The distribution of these two metabolites across the three sample locations matches well with the known enrichment of lutein and zeaxanthin in the macula. Furthermore, zeaxanthin is known to have a “sharper peak” than lutein in the fovea, when comparing macula versus periphery (**Figure S7**).

The metabolic missingness rate was 26% (sd=34%) among all samples with an increasing missingness rate between the areas (24% macula; 25.7% temporal, 28.4% periphery). Missingnessness in metabolomics study is usually assumed to arise from two different mechanisms, either Missingness At Random (MAR, the missingness of the metabolite does not depend on the value of the metabolite itself) or Missingness Not At Random (MNAR, the missingness of the metabolite depends on the value of the metabolite itself). Imputation of metabolic abundance depends on such the assumption of mechanism, however, no common consensus has yet been reached on what type should be assumed ^35–37^. Given the aforementioned result it is plausible to assume that some metabolites in this study have retinal region specific abundance and additionally present as MNAR. Which metabolites were responsible for this effect was, however, unclear. To this end, we performed imputation of metabolic abundances in a flexible manner by using two different imputation approaches depending on the type of missingness that was most likely for each metabolite. Firstly, we discarded from further analyses 62 metabolites (**Table S3**) with missingness rates of ≥80%. Secondly, for each metabolite *i* with at least one missing value, we identified two auxiliary metabolites (*Ji* and *Zi*) which had the two highest correlations with metabolite *i* (calculated over at least three observable abundances). The average correlation between metabolites and the first auxiliary variable J was r=0.92 (sd= 0.06) while a correlation of r=0.89 (sd= 0.07) was observed for the auxiliary variable

Z. Thirdly, auxiliary variables were then used for each metabolite to determine whether the missingness for that metabolite was MAR or MNAR. To this end, we performed a T-test comparing the observed values of both auxiliary variables when the metabolite of interest was missing or non-missing. We could not perform such a test for 41 metabolites as these presented with either one or two missing values. If metabolic values of either auxiliary variable were identified as being significantly lower (p<0.05) when metabolite *i* presented missing values then the missingness of metabolite *i* was considered MNAR. If neither of the two auxiliary variables were significantly lower or metabolite *i* had no auxiliary variables available it was considered to present as MAR. This resulted in 132 metabolites flagged as MAR and

37 as MNAR missingness. The metabolomics dataset was then imputed using the MetImp software v1.2 (https://metabolomics.cc.hawaii.edu/software/MetImp/) ^37,38^ using an MAR random forest approach which uses all available metabolic observations to impute missing values. Metabolites with MNAR missingness were imputed using the same tool with a Gibbs sampling approach. Both approaches have been used by several previous studies^35–37^.

We then proceeded to quantile normalise each sample to reduce the effect of potential confounding due to batch effect using the *NormaliseBetweenArrays* function of the Limma package v 3.44.3. We assessed the gain of biological separation between samples by comparing the first two metabolic principal components in the non-normalised data versus the normalised data (**Figure S8 A-B**). Investigation of preparation batches in the normalised data revealed an effect of batch preparation on metabolic abundance which was orthogonal to the biological differences **(Figure S8 C**).

Two metabolites were discarded due to the lack of any variability. The macula sample of one animal was discarded from further analyses since it was identified as incorrectly labelled based on clustering in the PC plots and examination of its carotene diol 1 and 3 levels. The final dataset comprised 32 samples and 371 metabolites. Metabolites were divided into 34 biological pathway groups. The list of metabolites and respective pathway membership is available in **Table S4**.

The data processing of the human serum metabolic data for the AMD study is presented in **Supplementary Methods** and are similar to those described above as well as a previously reported MacTel metabolomics study (Bonelli et al 2020).

## Statistical Analysis

To test for differential abundance between different areas of the retina we adopted the same strategy as in our previous MacTel metabolite study^5^. This strategy involved the usage of the Limma software for gene expression analysis which exploits multivariate linear regression combined with the empirical Bayes approach ^39–41^. Metabolites were average centred and standard deviation scaled. Primate age and gender, as well as preparation batch, were included as covariates in the model. Intra-primate correlation was taken into account by considering samples as biological replicates of the same primate using the function *duplicateCorrelation* in Limma ^42^. For each metabolite, we tested three contrasts: *Macular vs Temporal, Temporal vs Periphery* and *Macula vs Periphery*. For this study, we used a false discovery rate cut-off of 5%. All metabolites with a Benjamini–Hochberg corrected p-value less than the FDRcut off of 0.05 were considered significant.

To test for pathway differential abundance between retinal areas we tested for enrichment of differential abundance in the pathways by using the *fry* function from the R/Limma package. The reason behind using these two different approaches has been described elsewhere ^5^). Additionally, we prepared global pathway abundance by calculating the first principal component on all metabolites in each pathway. The specific methodology to perform this has been described elsewhere^5^

Metabolites and metabolic pathways were then divided into clusters reflecting the patterns of significance and effect direction between contrasts (*Macular vs Temporal, Temporal vs Periphery, Macula vs Periphery*). A table of cluster names with contrast combinations is presented in **Table S5**.

To assess the utility of our primate metabolomic study results for the interpretation of human retinal disease we extracted the results from our previous MacTel study ^5^ as well as the new study presented in this manuscript for AMD. Each metabolic family was divided into three subgroups dividing metabolites that were either, “enriched”, “not-differentiated” or “depleted” in the macula compared to the rest of the retina. With the new definition, we then tested for enrichment in each newly defined pathway using the R/limma *fry* module ^43^. Pathways with enrichment FDR<0.05 were considered significant in MacTel. Given the very low level of significance in the AMD study, a less stringent threshold of a nominal p-value < 0.05 was used instead for the enrichment analysis of this disorder.

Statistical analysis methodology used to analyse the serum metabolites of human samples comparing AMD to healthy controls is presented in **Supplementary Methods**.

