## Supplementary Materials for "Spatial distribution of metabolites in primate retina and its relevance to studies of human metabolic retinal disorders"

### Supplementary Methods

#### Data cleaning - Human Serum AMD vs Controls

Given their strongly skewed distribution, metabolic abundances were log-transformed to achieve distribution symmetry. Metabolic abundances were normalised by dividing the global median log-abundance of area/batch combination. Principal components plot revealed 3 subjects belonging to the same batch which were global outliers, these were discarded from further analyses (**Figure S9**).

Before imputation we discarded 209 metabolites presenting missingness rates higher than 20% in any of the 4 subgroups. Imputation was performed in the same fashion as in the primate study with 132 metabolites imputed assuming MCAR missingness and 110 with MNAR missingness. Quantile normalisation was performed as described in the primate study. No metabolites and no further samples were discarded after quantile normalization. The final dataset was composed by 348 samples and 763 metabolites. Metabolites were divided into 50 groups highlighting their biological pathways. The list of metabolites and relative groups is available in **Table S6**.

#### Statistical Analysis - Human Serum AMD vs Controls

Metabolic differential abundance was tested using the same approach as described for the primate study. In model contained in age, sex at birth and processing batch as covariates. No intra-subject correlation was taken into account since only one observation was included per subject for each metabolite. To achieve maximal discovery power, subject-specific weights were calculated using the function *arrayWeights* from the Limma package and were included in the models. We performed two main analyses, one testing for all AMD patients against controls and another one testing each sub disease again controls. Enrichment analysis was performed for both main analyses as previously described and no principal component analysis was performed for this study.

### Supplementary Results

#### Metabolite missingness in primate data

Given the relatively small number of detected metabolites and their strong correlations with each other, in this study, we used a somewhat forgiving missingness threshold to discard a-priority any metabolite (>80%). However, we found that only 44 significant metabolites had an initial missingness rate greater than 20% and only 23 greater than 50% (**Table S4**). We tested whether significance was higher for those metabolites presenting different missingness rates between areas as a quality check. We found that significant metabolites had an average standard deviation of missingness across areas lower than those non-significant (AveMissSD 0.055 vs 0.068) which confirmed that our results were not biased towards metabolites presenting different missingness rates across areas.

#### Metabolite differential abundance AMD vs Controls

We found a total of 4 metabolites whose abundance was significantly different between AMD cases and controls. All these were depleted and only significant when comparing CNV-AMD to controls and only 1 of these was significant when comparing all AMD cases against controls (**Table S6, Figure S10**).

#### Pathway differential abundance AMD vs Controls

We then investigated whether metabolites belonging to the same metabolic pathway shared a similar abundance difference across retinal areas. We found two biological pathways to be enriched with metabolites sharing similar signals for disease status. Both were only significant when comparing CNV-AMD to controls and only one of these was significant when comparing all AMD cases against controls (**Table S6**, **Figure S11**).

### Supplementary Figures

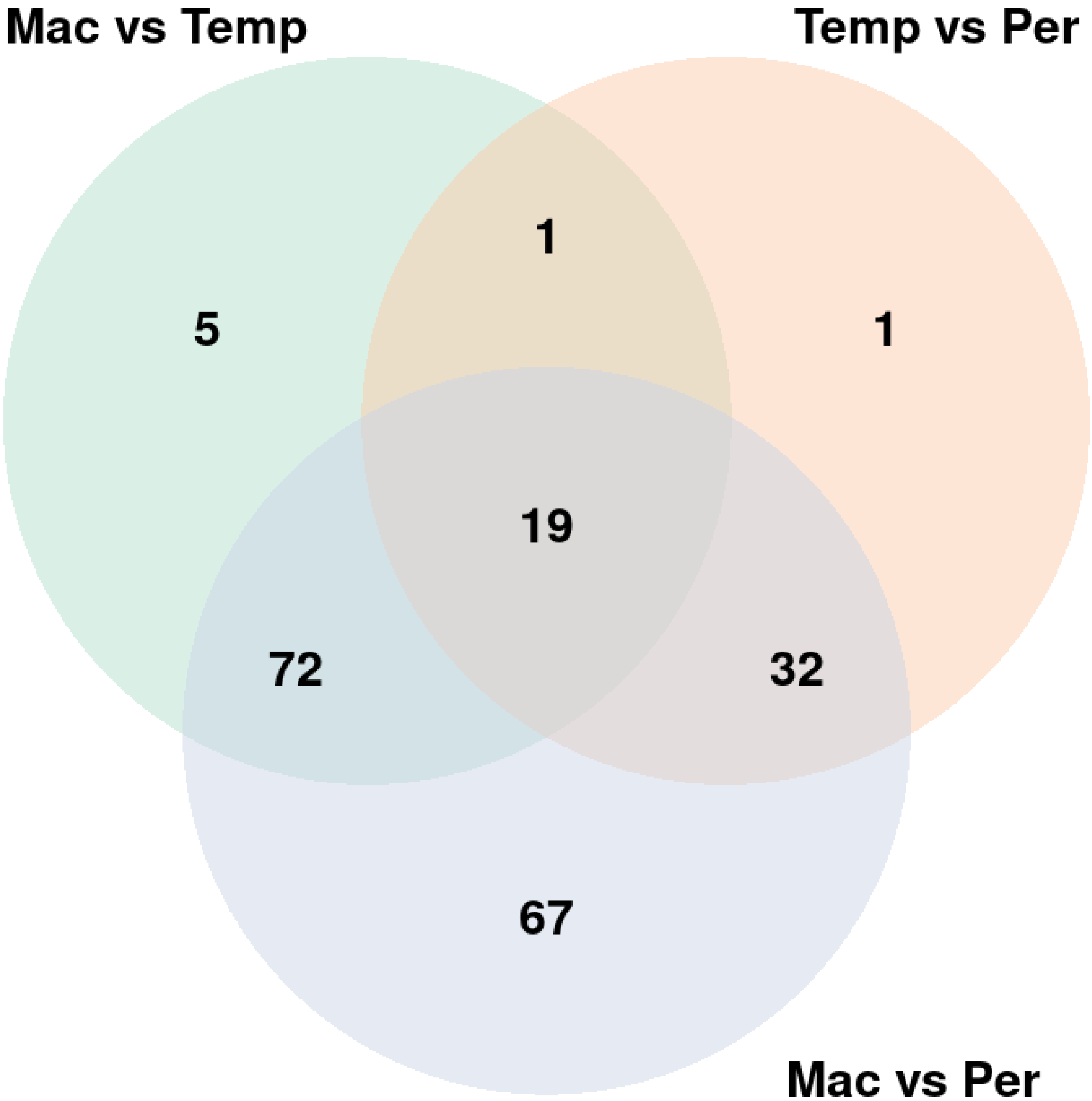


**Figure S1:** Venn diagram representing frequencies of significant metabolite by the three contrasts *Macula vs Temporal* (Mac vs Temp), *Temporal vs Periphery* (Temp vs Per), and *Macula vs Periphery* (Mac vs Per).

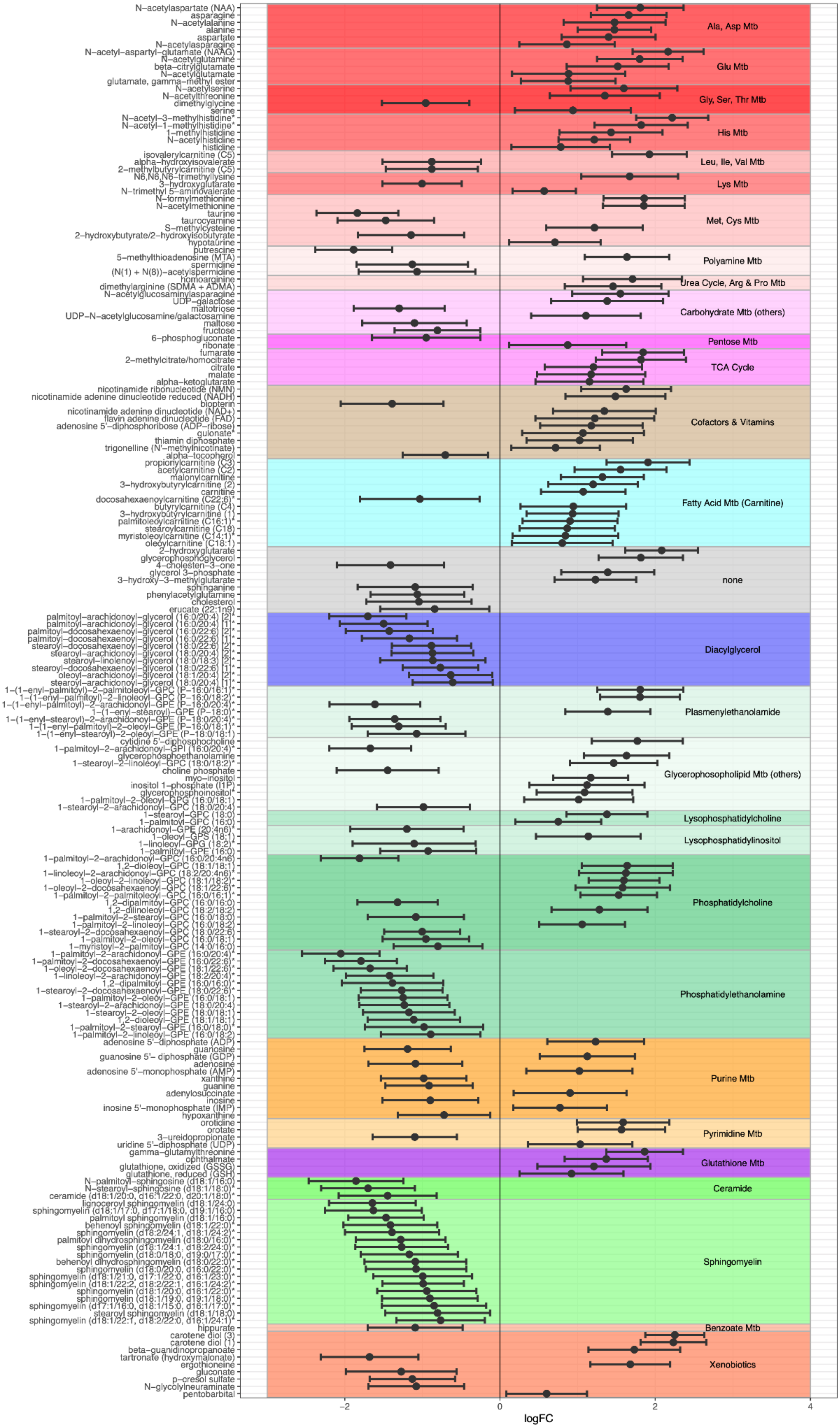

N–acetylaspartate (NAA)
asparagine
N–acetylalanine
alanine
aspartate
N–acetylasparagine
N–acetyl–aspartyl–glutamate (NAAG)
N–acetylglutamine
beta–citrylglutamate
N–acetylglutamate
glutamate, gamma–methyl ester
N–acetylserine
N–acetylthreonine
dimethylglycine
serine
N–acetyl–3–methylhistidine*
N–acetyl–1–methylhistidine*
1–methylhistidine
N–acetylhistidine
histidine
isovalerylcarnitine (C5)
alpha–hydroxyisovalerate
2–methylbutyrylcarnitine (C5)
N6,N6,N6–trimethyllysine
3–hydroxyglutarate
N–trimethyl 5–aminovalerate
N–formylmethionine
N–acetylmethionine
taurine
taurocyamine
S–methylcysteine
2–hydroxybutyrate/2–hydroxyisobutyrate
hypotaurine
putrescine
5–methylthioadenosine (MTA)
spermidine
(N(1) + N(8))–acetylspermidine
homoarginine
dimethylarginine (SDMA + ADMA)
N–acetylglucosaminylasparagine
UDP–galactose
maltotriose
UDP–N–acetylglucosamine/galactosamine
maltose
fructose
6–phosphogluconate
ribonate
fumarate
2–methylcitrate/homocitrate
citrate
malate
alpha–ketoglutarate
nicotinamide ribonucleotide (NMN)
nicotinamide adenine dinucleotide reduced (NADH)
biopterin
nicotinamide adenine dinucleotide (NAD+)
flavin adenine dinucleotide (FAD)
adenosine 5'–diphosphoribose (ADP–ribose)
gulonate*
thiamin diphosphate
trigonelline (N'–methylnicotinate)
alpha–tocopherol
propionylcarnitine (C3)
acetylcarnitine (C2)
malonylcarnitine
3–hydroxybutyrylcarnitine (2)
carnitine
docosahexaenoylcarnitine (C22:6)*
butyrylcarnitine (C4)
3–hydroxybutyrylcarnitine (1)
palmitoleoylcarnitine (C16:1)*
stearoylcarnitine (C18)
myristoleoylcarnitine (C14:1)*
oleoylcarnitine (C18:1)
2–hydroxyglutarate
glycerophosphoglycerol
4–cholesten–3–one
glycerol 3–phosphate
3–hydroxy–3–methylglutarate
sphinganine
phenylacetylglutamine
cholesterol
erucate (22:1n9)
palmitoyl–arachidonoyl–glycerol (16:0/20:4) [2]*
palmitoyl–arachidonoyl–glycerol (16:0/20:4) [1]*
palmitoyl–docosahexaenoyl–glycerol (16:0/22:6) [2]*
palmitoyl–docosahexaenoyl–glycerol (16:0/22:6) [1]*
stearoyl–docosahexaenoyl–glycerol (18:0/22:6) [2]*
stearoyl–arachidonoyl–glycerol (18:0/20:4) [2]*
stearoyl–linolenoyl–glycerol (18:0/18:3) [2]*
stearoyl–docosahexaenoyl–glycerol (18:0/22:6) [1]*
oleoyl–arachidonoyl–glycerol (18:1/20:4) [2]*
stearoyl–arachidonoyl–glycerol (18:0/20:4) [1]*
1–(1–enyl–palmitoyl)–2–palmitoleoyl–GPC (P–16:0/16:1)*
1–(1–enyl–palmitoyl)–2–linoleoyl–GPC (P–16:0/18:2)*
1–(1–enyl–palmitoyl)–2–arachidonoyl–GPE (P–16:0/20:4)*
1–(1–enyl–stearoyl)–GPE (P–18:0)*
1–(1–enyl–stearoyl)–2–arachidonoyl–GPE (P–18:0/20:4)*
1–(1–enyl–palmitoyl)–2–oleoyl–GPE (P–16:0/18:1)*
1–(1–enyl–stearoyl)–2–oleoyl–GPE (P–18:0/18:1)
cytidine 5'–diphosphocholine
1–palmitoyl–2–arachidonoyl–GPI (16:0/20:4)*
glycerophosphoethanolamine
1–stearoyl–2–linoleoyl–GPC (18:0/18:2)*
choline phosphate
myo–inositol
inositol 1–phosphate (I1P)
glycerophosphoinositol*
1–palmitoyl–2–oleoyl–GPG (16:0/18:1)
1–stearoyl–2–arachidonoyl–GPC (18:0/20:4)
1–stearoyl–GPC (18:0)
1–palmitoyl–GPC (16:0)
1–arachidonoyl–GPE (20:4n6)*
1–oleoyl–GPS (18:1)
1–linoleoyl–GPG (18:2)*
1–palmitoyl–GPE (16:0)
1–palmitoyl–2–arachidonoyl–GPC (16:0/20:4n6)
1,2–dioleoyl–GPC (18:1/18:1)
1–linoleoyl–2–arachidonoyl–GPC (18:2/20:4n6)*
1–oleoyl–2–linoleoyl–GPC (18:1/18:2)*
1–oleoyl–2–docosahexaenoyl–GPC (18:1/22:6)*
1–palmitoyl–2–palmitoleoyl–GPC (16:0/16:1)*
1,2–dipalmitoyl–GPC (16:0/16:0)
1,2–dilinoleoyl–GPC (18:2/18:2)
1–palmitoyl–2–stearoyl–GPC (16:0/18:0)
1–palmitoyl–2–linoleoyl–GPC (16:0/18:2)
1–stearoyl–2–docosahexaenoyl–GPC (18:0/22:6)
1–palmitoyl–2–oleoyl–GPC (16:0/18:1)
1–myristoyl–2–palmitoyl–GPC (14:0/16:0)
1–palmitoyl–2–arachidonoyl–GPE (16:0/20:4)*
1–palmitoyl–2–docosahexaenoyl–GPE (16:0/22:6)*
1–oleoyl–2–docosahexaenoyl–GPE (18:1/22:6)*
1–linoleoyl–2–arachidonoyl–GPE (18:2/20:4)*
1,2–dipalmitoyl–GPE (16:0/16:0)*
1–stearoyl–2–docosahexaenoyl–GPE (18:0/22:6)*
1–palmitoyl–2–oleoyl–GPE (16:0/18:1)
1–stearoyl–2–arachidonoyl–GPE (18:0/20:4)
1–stearoyl–2–oleoyl–GPE (18:0/18:1)
1,2–dioleoyl–GPE (18:1/18:1)
1–palmitoyl–2–stearoyl–GPE (16:0/18:0)*
1–palmitoyl–2–linoleoyl–GPE (16:0/18:2)
adenosine 5'–diphosphate (ADP)
guanosine
guanosine 5'– diphosphate (GDP)
adenosine
adenosine 5'–monophosphate (AMP)
xanthine
guanine
adenylosuccinate
inosine
inosine 5'–monophosphate (IMP)
hypoxanthine
orotidine
orotate
3–ureidopropionate
uridine 5'–diphosphate (UDP)
gamma–glutamylthreonine
ophthalmate
glutathione, oxidized (GSSG)
glutathione, reduced (GSH)
N–palmitoyl–sphingosine (d18:1/16:0)
N–stearoyl–sphingosine (d18:1/18:0)*
ceramide (d18:1/20:0, d16:1/22:0, d20:1/18:0)*
lignoceroyl sphingomyelin (d18:1/24:0)
sphingomyelin (d18:1/17:0, d17:1/18:0, d19:1/16:0)
palmitoyl sphingomyelin (d18:1/16:0)
behenoyl sphingomyelin (d18:1/22:0)*
sphingomyelin (d18:2/24:1, d18:1/24:2)*
palmitoyl dihydrosphingomyelin (d18:0/16:0)*
sphingomyelin (d18:1/24:1, d18:2/24:0)*
sphingomyelin (d18:0/18:0, d19:0/17:0)*
behenoyl dihydrosphingomyelin (d18:0/22:0)*
sphingomyelin (d18:0/20:0, d16:0/22:0)*
sphingomyelin (d18:1/21:0, d17:1/22:0, d16:1/23:0)*
sphingomyelin (d18:1/22:2, d18:2/22:1, d16:1/24:2)*
sphingomyelin (d18:1/20:0, d16:1/22:0)*
sphingomyelin (d18:1/19:0, d19:1/18:0)*
sphingomyelin (d17:1/16:0, d18:1/15:0, d16:1/17:0)*
stearoyl sphingomyelin (d18:1/18:0)
sphingomyelin (d18:1/22:1, d18:2/22:0, d16:1/24:1)*
hippurate
carotene diol (3)
carotene diol (1)
beta–guanidinopropanoate
tartronate (hydroxymalonate)
ergothioneine
gluconate
p–cresol sulfate
N–glycolylneuraminate
pentobarbital
Ala, Asp Mtb
Glu Mtb
Gly, Ser, Thr Mtb
His Mtb
Leu, Ile, Val Mtb
Lys Mtb
Met, Cys Mtb
Polyamine Mtb
Urea Cycle, Arg & Pro Mtb
Carbohydrate Mtb (others)
Pentose Mtb
TCA Cycle
Cofactors & Vitamins
Fatty Acid Mtb (Carnitine)
none
Diacylglycerol
Plasmenylethanolamide
Glycerophosopholipid Mtb (others)
Lysophosphatidylcholine
Lysophosphatidylinositol
Phosphatidylcholine
Phosphatidylethanolamine
Purine Mtb
Pyrimidine Mtb
Glutathione Mtb
Ceramide
Sphingomyelin
Benzoate Mtb
Xenobiotics
−2
0
2
4
logFC

**Figure S2:** Log-fold changes and 95% confidence interval of metabolites with significant differential abundance between macula and periphery. Positive log-fold changes values in this figure indicate that the metabolite abundance was higher in the macula while negative represent higher abundance in the periphery. Metabolites have been divided and coloured by their respective biological pathways.

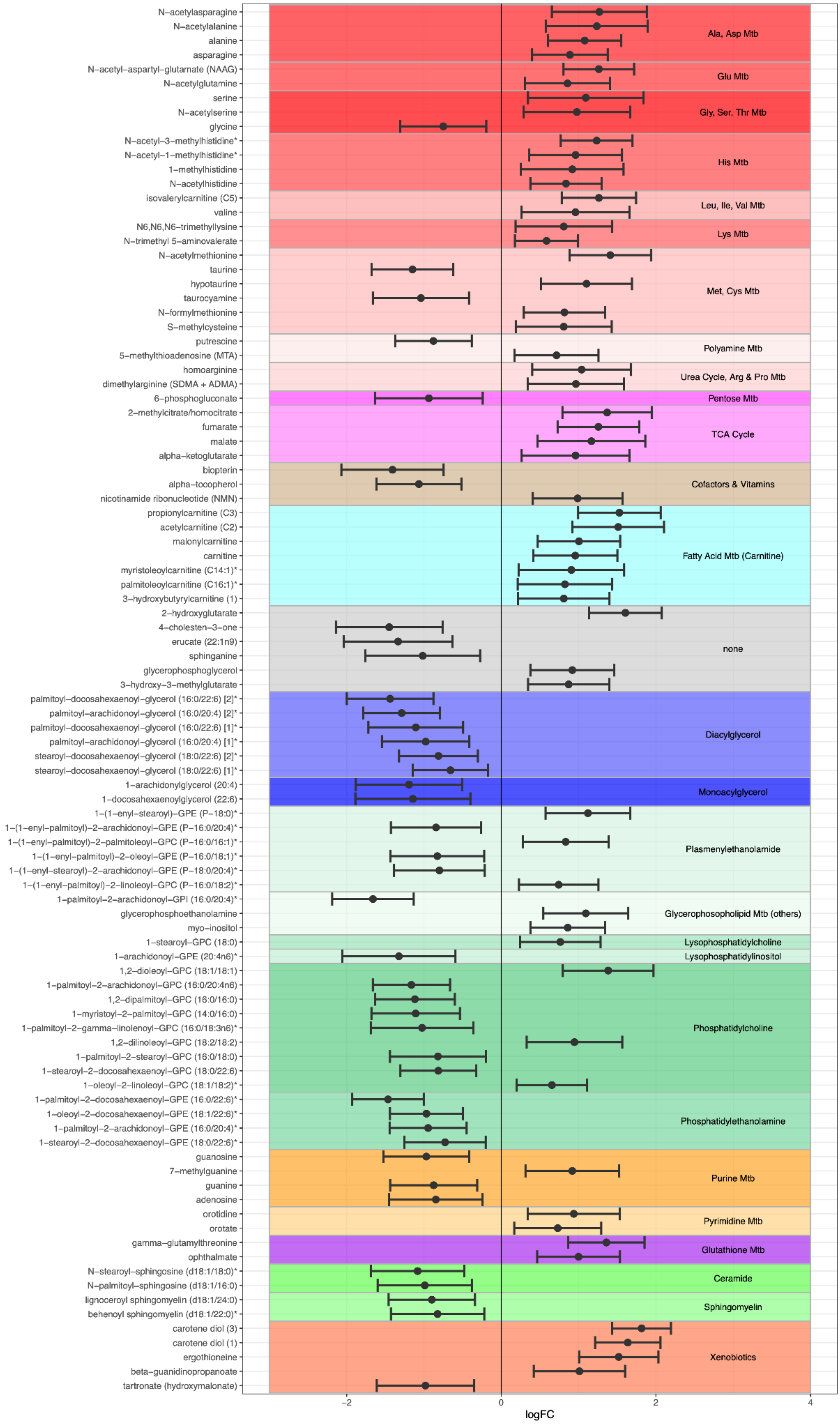

N–acetylasparagine
N–acetylalanine
alanine
asparagine
N–acetyl–aspartyl–glutamate (NAAG)
N–acetylglutamine
serine
N–acetylserine
glycine
N–acetyl–3–methylhistidine*
N–acetyl–1–methylhistidine*
1–methylhistidine
N–acetylhistidine
isovalerylcarnitine (C5)
valine
N6,N6,N6–trimethyllysine
N–trimethyl 5–aminovalerate
N–acetylmethionine
taurine
hypotaurine
taurocyamine
N–formylmethionine
S–methylcysteine
putrescine
5–methylthioadenosine (MTA)
homoarginine
dimethylarginine (SDMA + ADMA)
6–phosphogluconate
2–methylcitrate/homocitrate
fumarate
malate
alpha–ketoglutarate
biopterin
alpha–tocopherol
nicotinamide ribonucleotide (NMN)
propionylcarnitine (C3)
acetylcarnitine (C2)
malonylcarnitine
carnitine
myristoleoylcarnitine (C14:1)*
palmitoleoylcarnitine (C16:1)*
3–hydroxybutyrylcarnitine (1)
2–hydroxyglutarate
4–cholesten–3–one
erucate (22:1n9)
sphinganine
glycerophosphoglycerol
3–hydroxy–3–methylglutarate
palmitoyl–docosahexaenoyl–glycerol (16:0/22:6) [2]*
palmitoyl–arachidonoyl–glycerol (16:0/20:4) [2]*
palmitoyl–docosahexaenoyl–glycerol (16:0/22:6) [1]*
palmitoyl–arachidonoyl–glycerol (16:0/20:4) [1]*
stearoyl–docosahexaenoyl–glycerol (18:0/22:6) [2]*
stearoyl–docosahexaenoyl–glycerol (18:0/22:6) [1]*
1–arachidonylglycerol (20:4)
1–docosahexaenoylglycerol (22:6)
1–(1–enyl–stearoyl)–GPE (P–18:0)*
1–(1–enyl–palmitoyl)–2–arachidonoyl–GPE (P–16:0/20:4)*
1–(1–enyl–palmitoyl)–2–palmitoleoyl–GPC (P–16:0/16:1)*
1–(1–enyl–palmitoyl)–2–oleoyl–GPE (P–16:0/18:1)*
1–(1–enyl–stearoyl)–2–arachidonoyl–GPE (P–18:0/20:4)*
1–(1–enyl–palmitoyl)–2–linoleoyl–GPC (P–16:0/18:2)*
1–palmitoyl–2–arachidonoyl–GPI (16:0/20:4)*
glycerophosphoethanolamine
myo–inositol
1–stearoyl–GPC (18:0)
1–arachidonoyl–GPE (20:4n6)*
1,2–dioleoyl–GPC (18:1/18:1)
1–palmitoyl–2–arachidonoyl–GPC (16:0/20:4n6)
1,2–dipalmitoyl–GPC (16:0/16:0)
1–myristoyl–2–palmitoyl–GPC (14:0/16:0)
1–palmitoyl–2–gamma–linolenoyl–GPC (16:0/18:3n6)*
1,2–dilinoleoyl–GPC (18:2/18:2)
1–palmitoyl–2–stearoyl–GPC (16:0/18:0)
1–stearoyl–2–docosahexaenoyl–GPC (18:0/22:6)
1–oleoyl–2–linoleoyl–GPC (18:1/18:2)*
1–palmitoyl–2–docosahexaenoyl–GPE (16:0/22:6)*
1–oleoyl–2–docosahexaenoyl–GPE (18:1/22:6)*
1–palmitoyl–2–arachidonoyl–GPE (16:0/20:4)*
1–stearoyl–2–docosahexaenoyl–GPE (18:0/22:6)*
guanosine
7–methylguanine
guanine
adenosine
orotidine
orotate
gamma–glutamylthreonine
ophthalmate
N–stearoyl–sphingosine (d18:1/18:0)*
N–palmitoyl–sphingosine (d18:1/16:0)
lignoceroyl sphingomyelin (d18:1/24:0)
behenoyl sphingomyelin (d18:1/22:0)*
carotene diol (3)
carotene diol (1)
ergothioneine
beta–guanidinopropanoate
tartronate (hydroxymalonate)
Ala, Asp Mtb
Glu Mtb
Gly, Ser, Thr Mtb
His Mtb
Leu, Ile, Val Mtb
Lys Mtb
Met, Cys Mtb
Polyamine Mtb
Urea Cycle, Arg & Pro Mtb
Pentose Mtb
TCA Cycle
Cofactors & Vitamins
Fatty Acid Mtb (Carnitine)
none
Diacylglycerol
Monoacylglycerol
Plasmenylethanolamide
Glycerophosophoplipid Mtb (others)
Lysophosphatidylcholine
Lysophosphatidylinositol
Phosphatidylcholine
Phosphatidylethanolamine
Purine Mtb
Pyrimidine Mtb
Glutathione Mtb
Ceramide
Sphingomyelin
Xenobiotics
−2
0
2
4
logFC

**Figure S3:** Log-fold changes and 95% confidence interval of metabolites with significant differential abundance between macula and termporal. Positive log-fold changes values in this figure indicate that the metabolite abundance was higher in the macula while negative represent higher abundance in the periphery. Metabolites have been divided and coloured by their respective biological pathways.

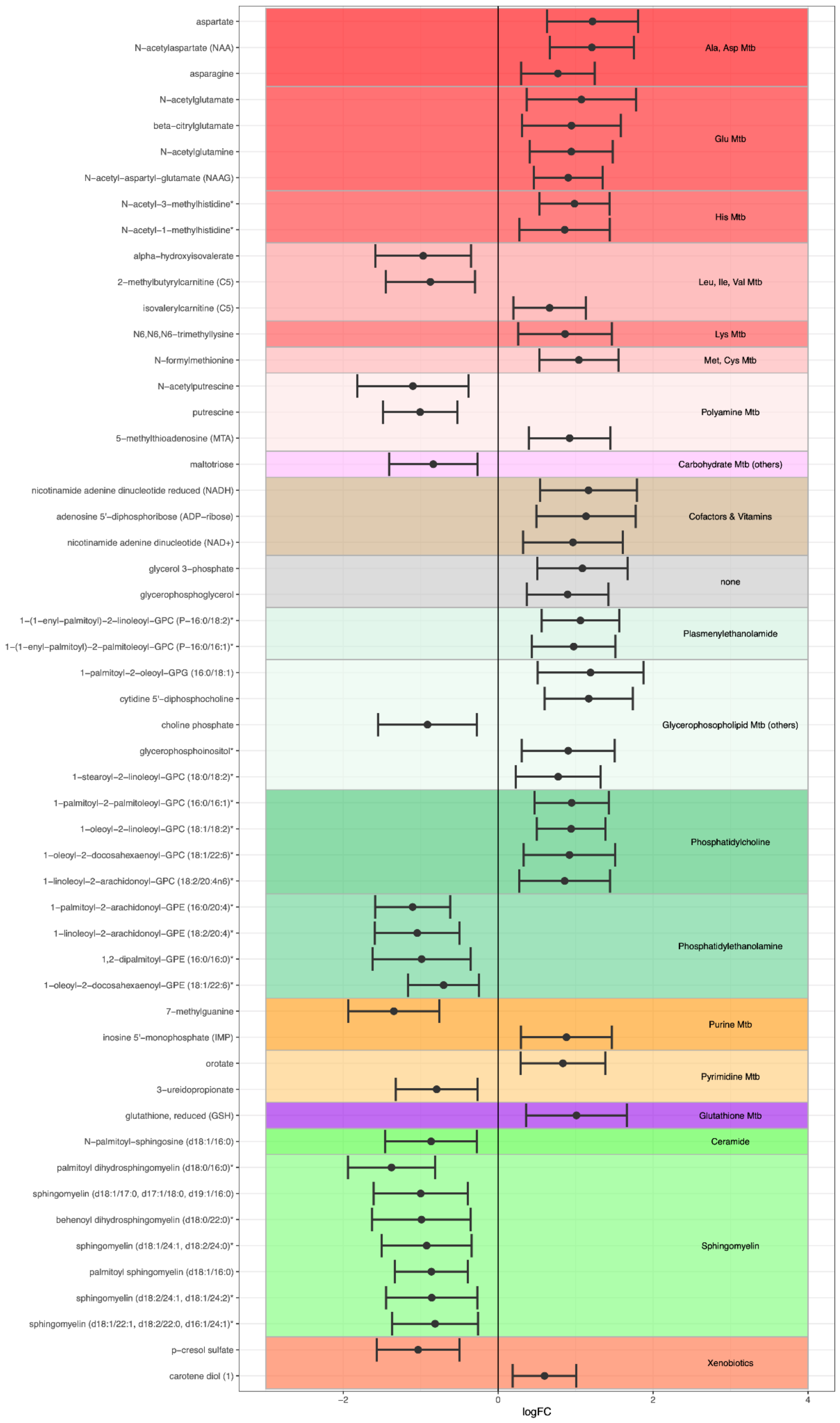

aspartate
N−acetylaspartate (NAA)
asparagine
N−acetylglutamate
beta−citrylglutamate
N−acetylglutamine
N−acetyl−aspartyl−glutamate (NAAG)
N−acetyl−3−methylhistidine*
N−acetyl−1−methylhistidine*
alpha−hydroxyisovalerate
2−methylbutyrylcarnitine (C5)
isovalerylcarnitine (C5)
N6,N6,N6−trimethyllysine
N−formylmethionine
N−acetylputrescine
putrescine
5−methylthioadenosine (MTA)
maltotriose
nicotinamide adenine dinucleotide reduced (NADH)
adenosine 5'−diphosphoribose (ADP−ribose)
nicotinamide adenine dinucleotide (NAD+)
glycerol 3−phosphate
glycerophosphoglycerol
1−(1−enyl−palmitoyl)−2−linoleoyl−GPC (P−16:0/18:2)*
1−(1−enyl−palmitoyl)−2−palmitoleoyl−GPC (P−16:0/16:1)*
1−palmitoyl−2−oleoyl−GPG (16:0/18:1)
cytidine 5'−diphosphocholine
choline phosphate
glycerophosphoinositol*
1−stearoyl−2−linoleoyl−GPC (18:0/18:2)*
1−palmitoyl−2−palmitoleoyl−GPC (16:0/16:1)*
1−oleoyl−2−linoleoyl−GPC (18:1/18:2)*
1−oleoyl−2−docosahexaenoyl−GPC (18:1/22:6)*
1−linoleoyl−2−arachidonoyl−GPC (18:2/20:4n6)*
1−palmitoyl−2−arachidonoyl−GPE (16:0/20:4)*
1−linoleoyl−2−arachidonoyl−GPE (18:2/20:4)*
1,2−dipalmitoyl−GPE (16:0/16:0)*
1−oleoyl−2−docosahexaenoyl−GPE (18:1/22:6)*
7−methylguanine
inosine 5'−monophosphate (IMP)
orotate
3−ureidopropionate
glutathione, reduced (GSH)
N−palmitoyl−sphingosine (d18:1/16:0)
palmitoyl dihydrosphingomyelin (d18:0/16:0)*
sphingomyelin (d18:1/17:0, d17:1/18:0, d19:1/16:0)
behenoyl dihydrosphingomyelin (d18:0/22:0)*
sphingomyelin (d18:1/24:1, d18:2/24:0)*
palmitoyl sphingomyelin (d18:1/16:0)
sphingomyelin (d18:2/24:1, d18:1/24:2)*
sphingomyelin (d18:1/22:1, d18:2/22:0, d16:1/24:1)*
p−cresol sulfate
carotene diol (1)
Ala, Asp Mtb
Glu Mtb
His Mtb
Leu, Ile, Val Mtb
Lys Mtb
Met, Cys Mtb
Polyamine Mtb
Carbohydrate Mtb (others)
Cofactors & Vitamins
none
Plasmenylethanolamide
Glycerophosopholipid Mtb (others)
Phosphatidylcholine
Phosphatidylethanolamine
Purine Mtb
Pyrimidine Mtb
Glutathione Mtb
Ceramide
Sphingomyelin
Xenobiotics
−2
0
2
4
logFC

**Figure S4:** Log-fold changes and 95% confidence interval of metabolites with significant differential abundance between temporal and periphery. Positive log-fold changes values in this figure indicate that the metabolite abundance was higher in the macula while negative represent higher abundance in the periphery. Metabolites have been divided and coloured by their respective biological pathways.

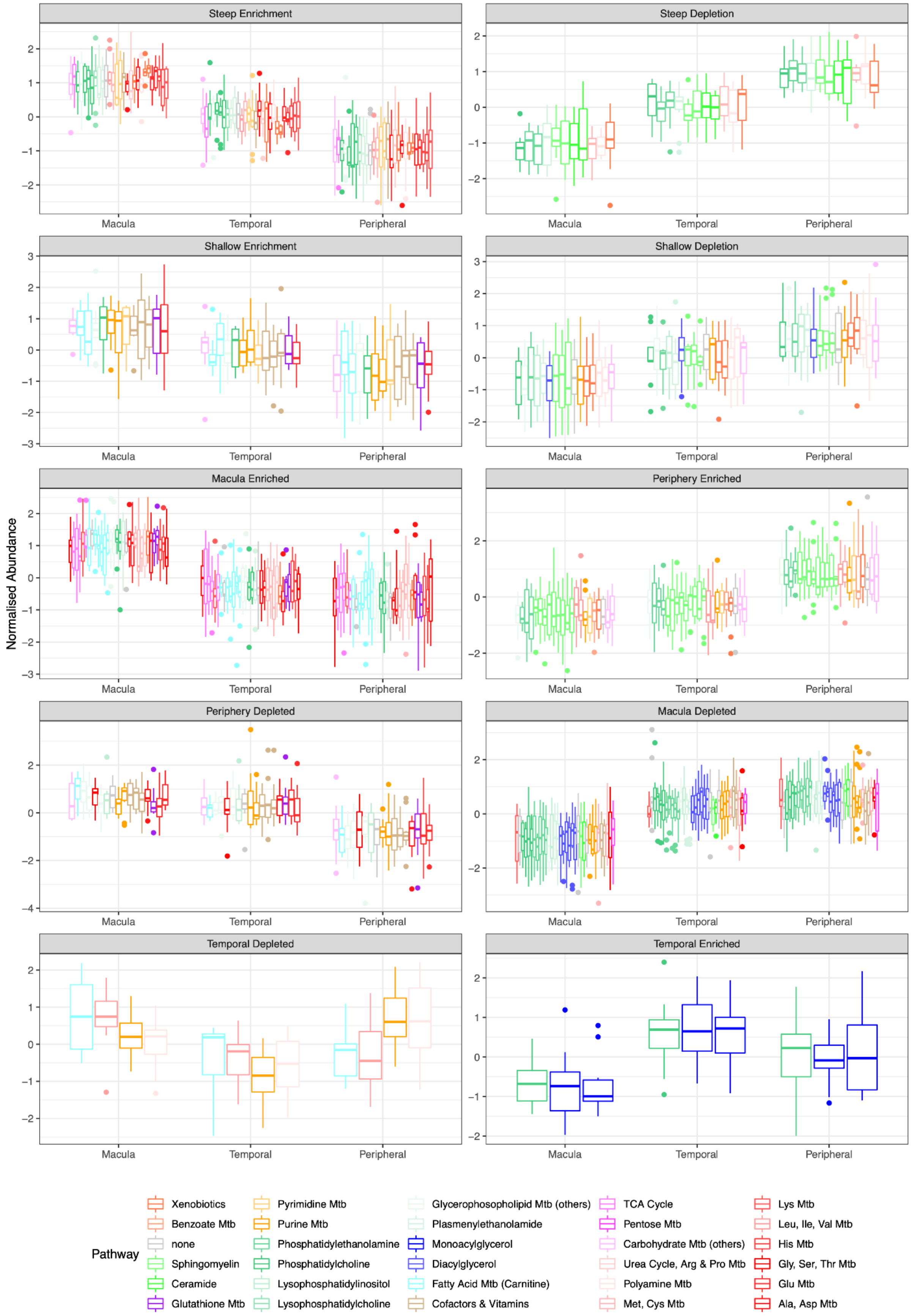


**Figure S5:** Metabolic distribution between clusters. In this image, every boxplot represents the covariate-corrected distribution of one metabolite whose abundance was significantly different for any of the three contrasts. Metabolites are divided into clusters according to their combination of significant results and respective log-fold change direction, further defined in Table 1 (summary statistics provided in **Table S4**). Boxplots have been coloured by the biological pathway of each metabolite. Increasing here is used to represent an increment of metabolic abundance from the macula to the periphery while decreasing represents the opposite.

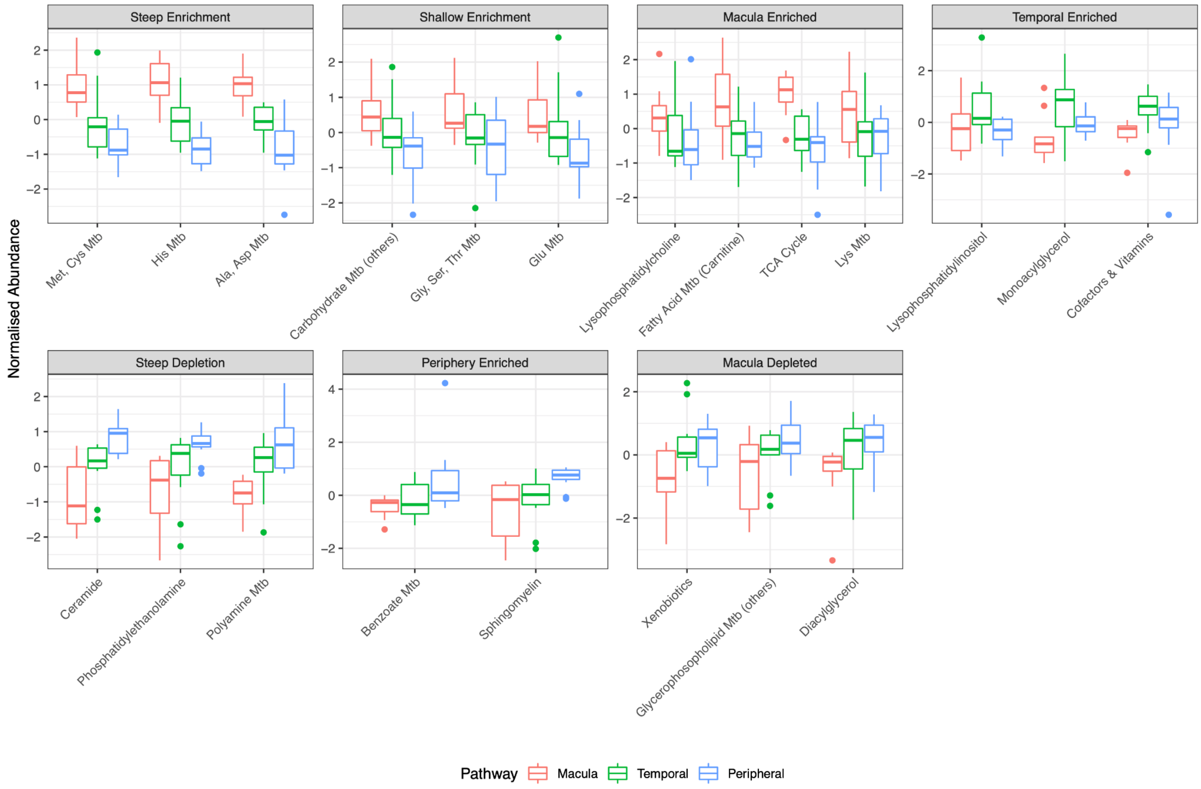


**Figure S6:** Pathway first PC distribution between retinal areas and divided by cluster. In this image, every boxplot represents the covariate-corrected distribution of the first PC for one pathway where this was significantly different for any of the three contrasts. Pathways are divided according to their combination of significant results and respective log-fold change direction (**Table S4**).

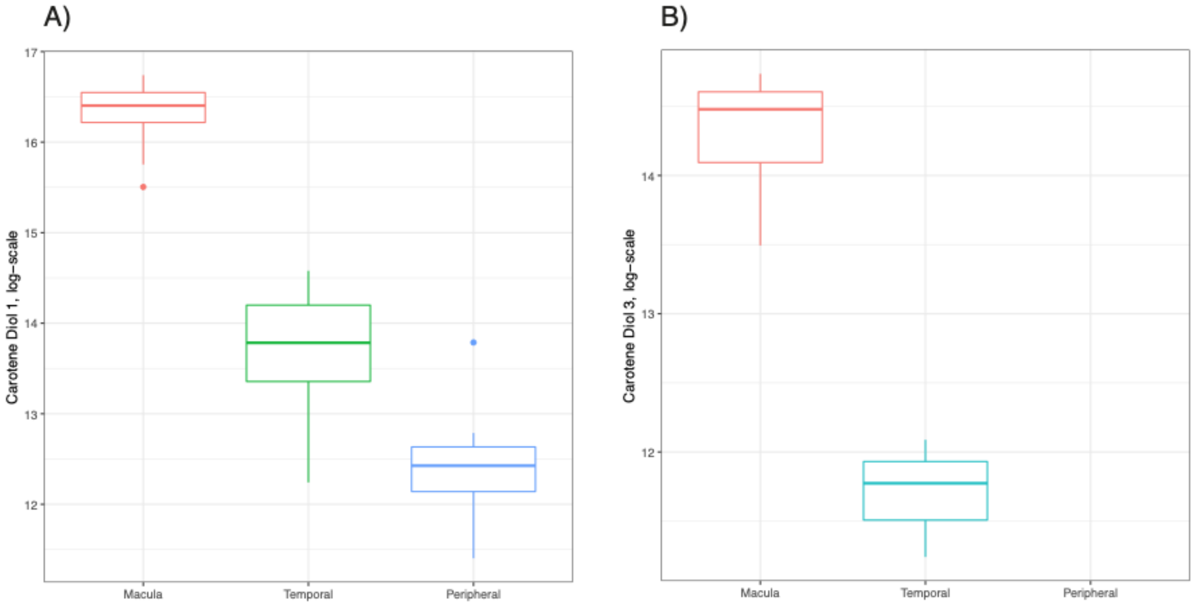


**Figure S7**: Positive control plot showing differences in (un-non-normalise, unnon-imputed) and log-transformed values of two carotenoids (most likely lutein and zeaxanthin) measured across samples collected across different retinal locations. A) Carotene diol 1 B) Carotene diol 3. Note the missing boxplot for carotene diol 3 in the peripheral area is due to complete missingness (i.e., below limit of machine detection) of this metabolite (i.e., below limit of machine detection) inobserved across peripheral samples.

A)

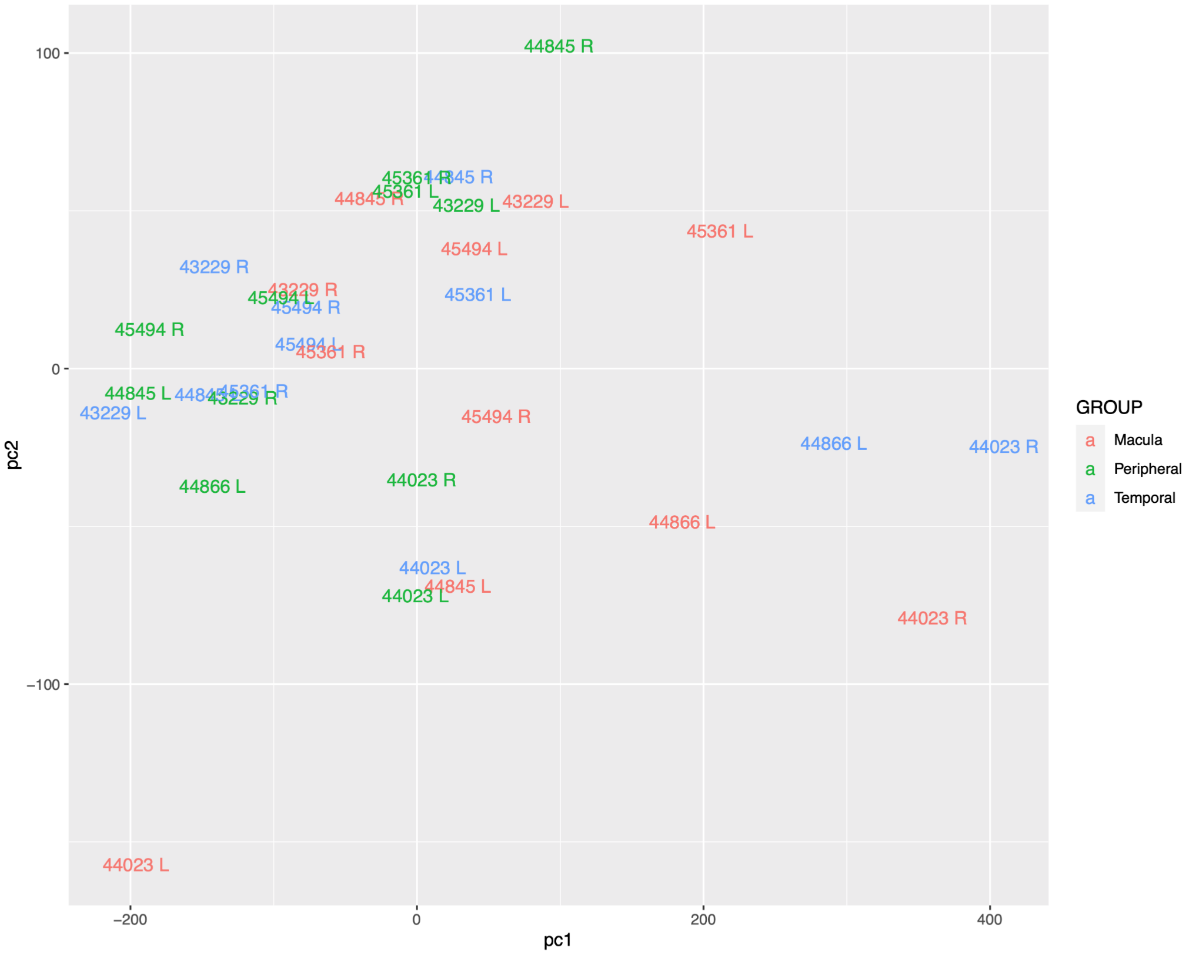
44845 R
45361 R
44845 R
45361 L
44845 R
43229 L
43229 L
45361 L
45494 L
43229 R
43229 R
45494 L
45494 R
45361 L
45494 R
45494 L
45361 R
44845 L
45361 R
43229 R
43229 L
45494 R
44866 L
44023 R
44023 R
44866 L
44023 L
44845 L
44023 L
44023 R
44023 L
100
0
−100
−200
200
400
pc1
pc2
GROUP
Macula
Peripheral
Temporal

B)

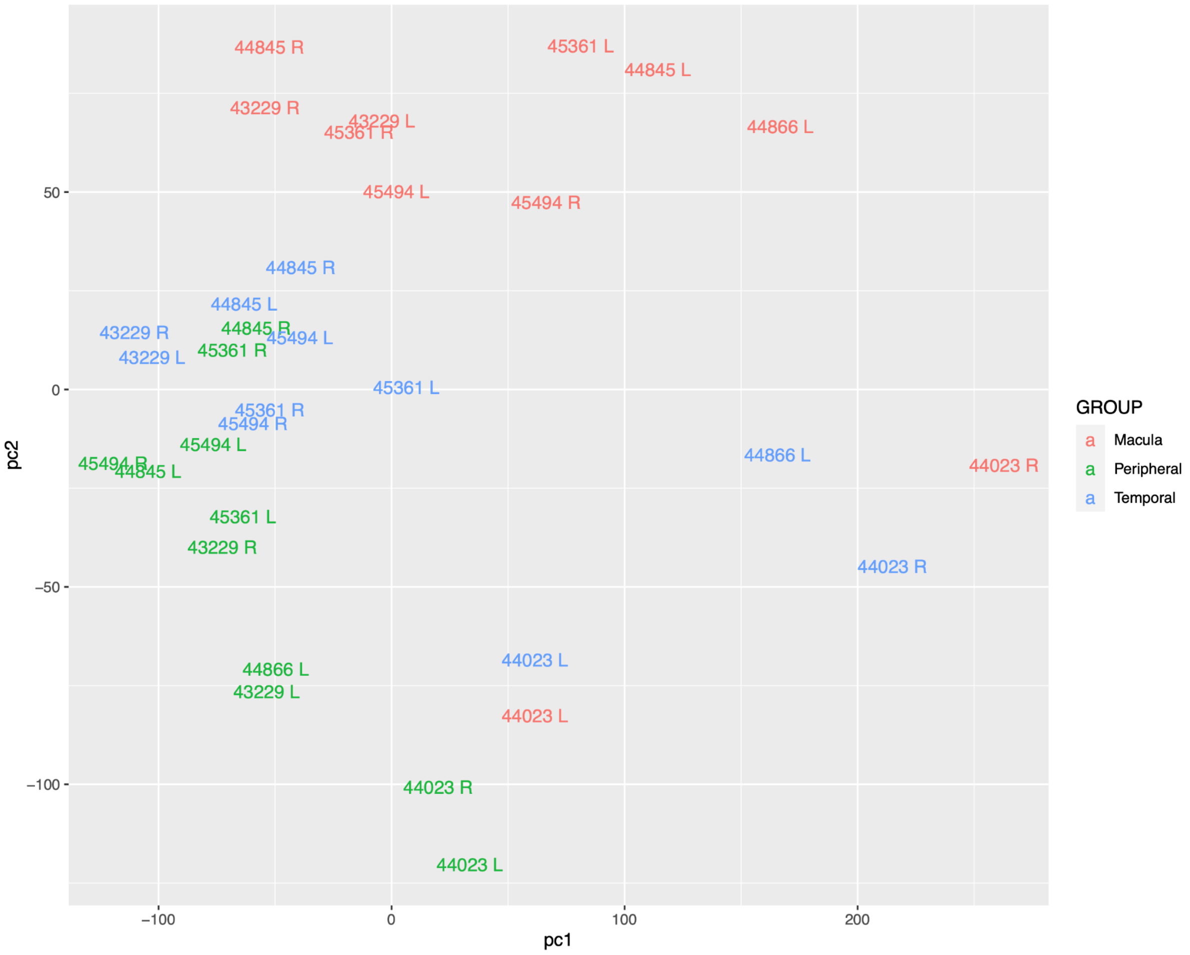
44845 R
45361 L
44845 L
43229 R
43229 L
45361 R
44866 L
45494 L
45494 R
44845 R
44845 L
43229 R
44845 R
45494 L
45361 R
43229 L
45361 L
45361 R
45494 R
45494 L
45494 R
44845 L
44866 L
44023 R
45361 L
43229 R
44023 R
44023 L
44866 L
43229 L
44023 L
44023 R
44023 L
50
0
−50
−100
−100
0
100
200
pc1
pc2
GROUP
Macula
Peripheral
Temporal

C)

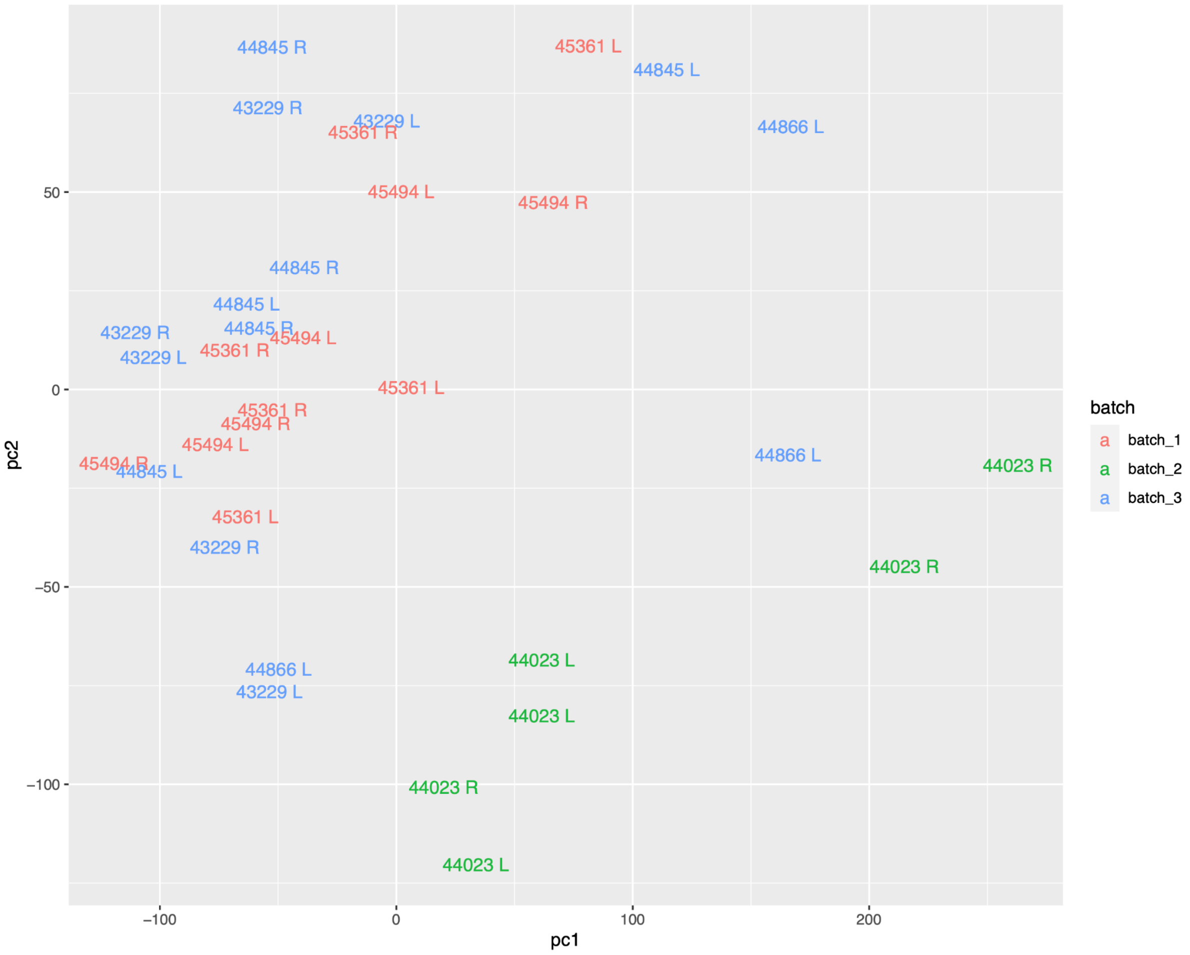


**Figure S8**: First two principal components plot showing the degrees of separation between biological samples in A) non-normalised data B) quantile normalised data. Animal ID number and eye (R=right eye, L=Left eye) is presented as text in the plot. C) First two principal components plot showing the degrees of separation between batches in the quantile normalised data.

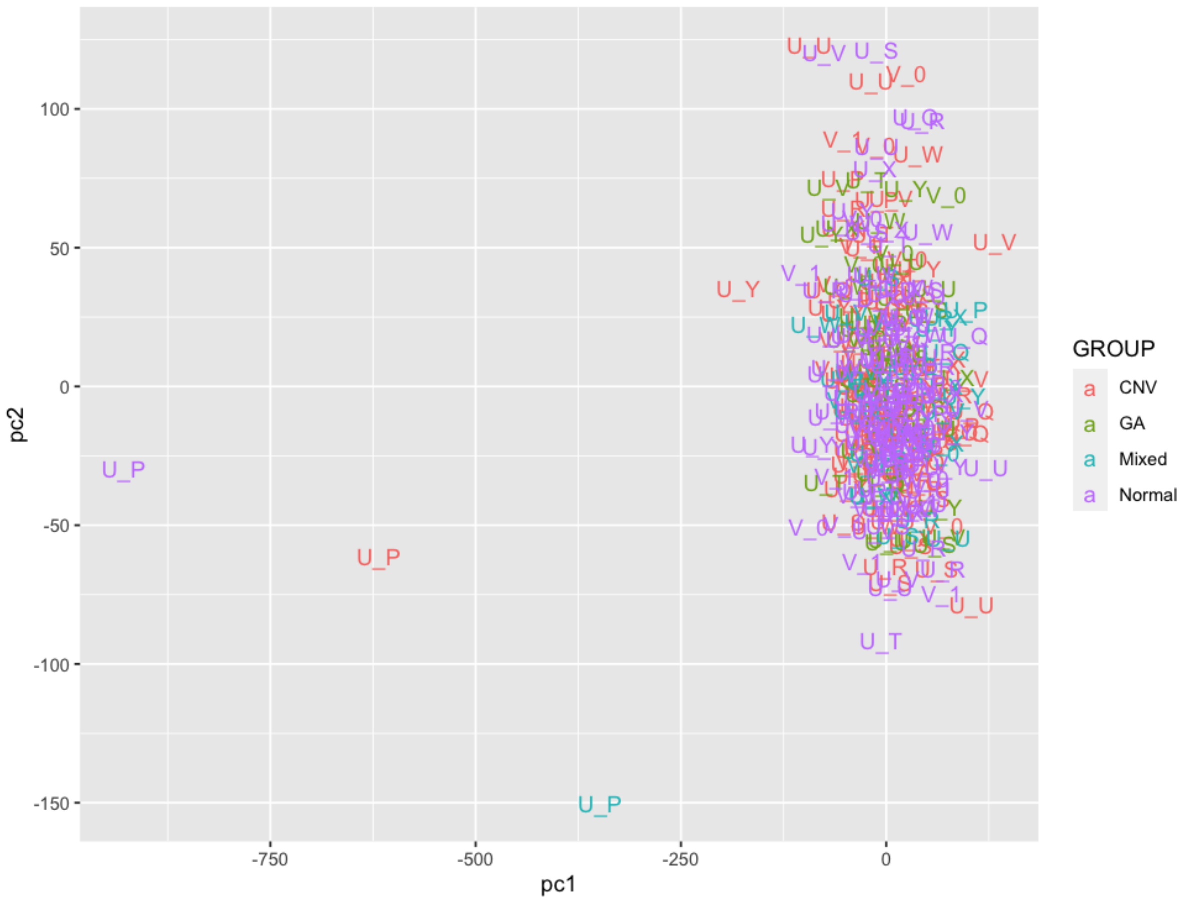


**Figure S9:** First two principal components plot on AMD metabolic data identifying 3 outliers. This plot every sample was colored accondying the the disease status and the string represented the sample preparation sub-bacth.

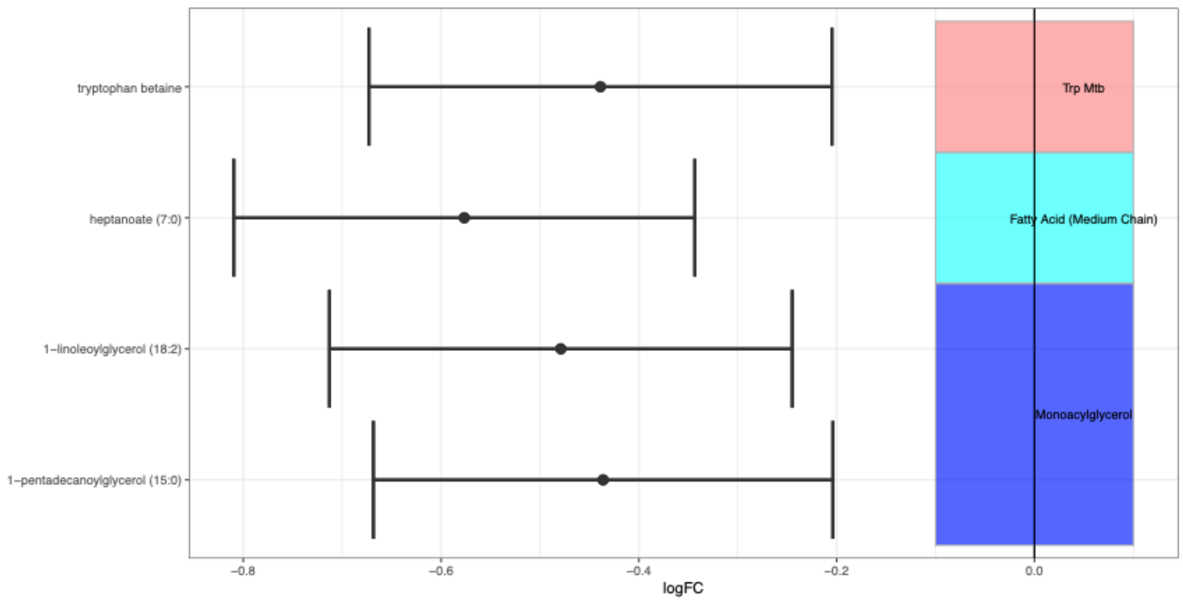


**Figure S10:** Log-fold changes and 95% confidence interval of metabolites with significant (FDR<0.05) differential abundance between CNV-AMD and healthy controls. Negative log-fold changes values in this figure indicate that the metabolite abundance was lower in CNV-AMD samples compared to controls. Metabolites have been divided and coloured by their respective biological pathways.

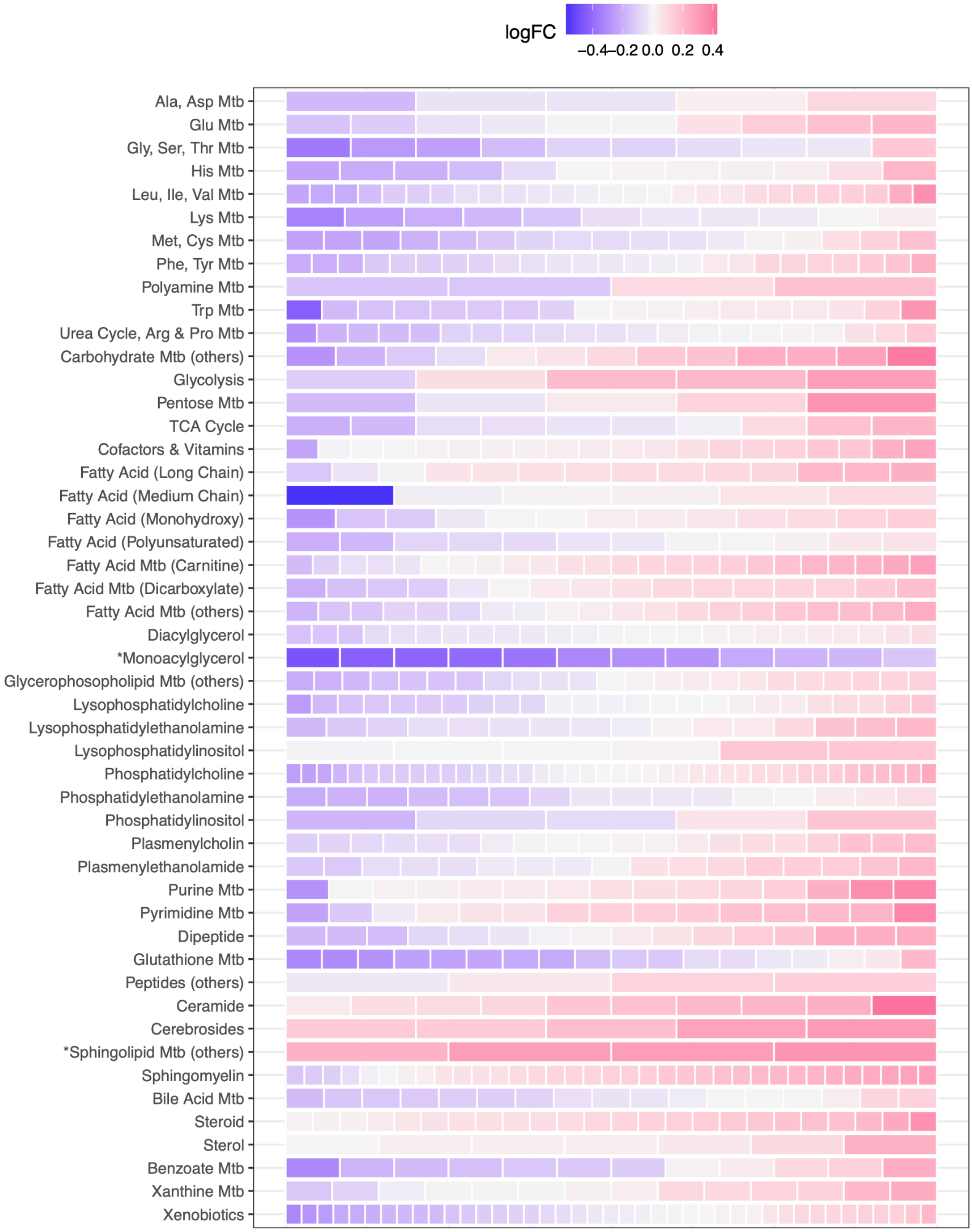


**Figure S11:** Pathways enrichment results. In this figure, each row represents a biological pathway. Metabolites are presented as “bricks” in each row. The colour of each metabolite represents the log-fold changes of that metabolite between CNV-AMD and healthy controls. High log-fold change indicates that the metabolite is more abundant in the CNV-AMD while negative values indicated that the metabolite is depleted. The symbol ‘*’ next to the pathway name indicates that that pathway was significantly enriched with either abundant or depleted metabolites in CNV-AMD compared to healthy controls.

### Supplementary Table Legends

**Table S1:** Table of differential abundance results for the primate analysis. This table contains most of the results for each metabolite presented in this publication. The results are divided into different categories: Differential Abundance By Metabolite, Pathway Enrichment Analysis, Missingness and Imputation, Additional Metabolites Info, and Pathway First PC Analysis results. (Note that a legend for each column presented in **Table S1** is provided as well as an enrichment analysis only version).
**Table S2:** Table of differential abundance results for the human serum AMD analysis. The results are divided into different categories: Differential Abundance By Metabolite, Pathway Enrichment Analysis, Missingness and Imputation, Additional Metabolites Info. (Note that a legend for each column presented in **Table S2** is provided).
**Table S3:** Table presenting the list of discarded metabolites. For each metabolite, a reason is presented for discarding it.
**Tabl2 S4:** Retained metabolites used in this study. This list contains all metabolites as well their classification in metabolic pathways as super-pathways.
**Table S5:** Table presenting metabolic clustering decision rules. For each cluster, the combination of significant effects for each contrast and their direction is presented.
